# Dietary supplementation with PUFAs rescues the eggshell defects caused by *seipin* mutations in *C. elegans*

**DOI:** 10.1101/2020.01.23.916718

**Authors:** Xiaofei Bai, Leng-Jie Huang, Sheng-Wen Chen, Ben Nebenfuehr, Brian Wysolmerski, Jui-Ching Wu, Sara K. Olson, Andy Golden, Chao-Wen Wang

## Abstract

SEIPIN, an ER membrane protein, plays critical roles in lipid droplet (LD) formation and lipid storage. Dysfunction of SEIPIN causes a variety of human diseases, including lipodystrophy, neuropathies, and male and female infertility. However, the cellular and molecular mechanisms of SEIPIN in causing these diseases are poorly understood. To address such mechanisms, we investigated the functional roles of *R01B10.6 (seip-1)*, the sole *SEIPIN1* ortholog in *C. elegans,* using CRISPR/Cas9 gene editing, and transcriptional assays. SEIP-1::mScarlet is widely expressed throughout development in *C. elegans*. Three full gene deletion mutants, generated by CRISPR/Cas9, displayed penetrant embryonic lethality. EM imaging and the visualization of reporter genes revealed that the lipid-rich permeability barrier, the innermost layer of the *C. elegans* embryonic eggshell, was defective or missing. Intriguingly, depletion of SEIP-1 revealed a perturbed gene expression pattern for fatty acid biosynthesis enzymes, in agreement with the disrupted permeability barrier formation phenotype of the embryos. Lastly, dietary supplementation of PUFAs rescued the embryonic lethality and defective permeability barrier in the deletion mutants. In sum, our study suggests that SEIP-1 may maternally regulate LD biogenesis and maintain lipid homeostasis to orchestrate the formation of the lipid-rich permeability barrier, which is crucial for eggshell formation and embryogenesis.

## Introduction

Lipid droplets (LDs) are ubiquitous cellular organelles for neutral lipid storage. The ER-localized protein SEIPIN, also known as BSCL2 in humans, has been implicated as a key player in LD biology (Cartwright and Goodman, 2012; Fei et al., 2011). Human *SEIPIN1* is highly expressed in brain, testis and adipose tissue (Magre et al., 2001). Missense mutations in *SEIPIN1* were identified in patients diagnosed with Beradinelli-Seip congenital generalized lipodystrophy type 2 (BSCL2) syndrome, motor neuropathy and Silver syndrome, suggesting the importance of SEIPIN1 for human lipid metabolism in different tissues (Chen et al., 2009; Fei et al., 2011; Ito et al., 2008; Magre et al., 2001; Walther et al., 2017; Windpassinger et al., 2004). Interestingly, a pair of compound *SEIPIN1* mutations were linked to teratozoospermia syndrome, which displays defective sperm and male infertility (Jiang et al., 2014). Consistent with these clinical studies, earlier investigations in mammalian and yeast cells have shown that loss-of-function SEIPIN causes defective fatty acid biogenesis and ectopic LD morphology, including tiny aggregated LDs and enlarged LDs (Fei et al., 2008; Grippa et al., 2015; Szymanski et al., 2007; Wang et al., 2016a; Wolinski et al., 2011). In mice, SEIPIN1 deficiency also leads to defective reproduction and infertility (El Zowalaty et al., 2015; Zowalaty and Ye, 2017). However, the cellular and molecular mechanisms of the *SEIPIN1*-associated genetic diseases remain largely unknown.

*Caenorhabditis elegans* is an attractive model for addressing fundamental questions about the metabolism of LDs. Although *C. elegans* lacks adipose tissue, such as adipocytes in mammals, they do store fats within abundant LDs, with TAGs being the major storage lipid, in the intestinal and hypodermal cells (Mak, 2012). Recent evidence suggests that LDs also exist in *C. elegans* oocytes and embryos (Shi et al., 2016). Unlike mammalian LDs, which measure up to 100 µm, *C. elegans* contains smaller LDs limited to 1-1.5 µm (Shi et al., 2013). Several LD-localized proteins have been characterized in *C. elegans*, such as triacylglycerol synthesis enzyme DGAT-2, fatty acid CoA synthetase ACS-22, and a major LD protein PLIN-1 that contributes to LD biogenesis (Golden et al., 2009; Li et al., 2016; Shi et al., 2013; Vrablik et al., 2015; Xu et al., 2012). Genetic mutants in *daf-2* IGF receptor and *daf-22* have been identified that result in aberrant LDs (Li et al., 2016). A recent study in *C. elegans* demonstrated that a deletion allele of *seip-1* reduced the size of a subset of LDs while over-expression had the opposite effect. This study also suggested that SEIP-1 was enriched in ER subdomains, called peri-LDs, and that this localization was dependent on polyunsaturated fatty acids (PUFAs) (Cao et al., 2019). Besides LDs, another lipid storage organelle, the yolk granule, is synthesized in the *C. elegans* intestine and transferred to maturing oocytes during reproduction (Hall et al., 1999; Kimble and Sharrock, 1983). This is similar to mammalian lipoproteins, which are synthesized in the liver and delivered to the ovum (Schneider, 1996). However, it is unclear whether the biosynthesis of yolk and LDs are related in *C. elegans*, and it is also unknown how LDs are generated and transported into the germ line, or whether they are generated *de novo* in oocytes. The yolk vitellogenin protein VIT-2, which is involved in the transfer of dietary lipids to LDs, may contribute to LD formation and regulation during reproduction (Vrablik et al., 2015; Wang et al., 2016b; Zhang et al., 2012). Failure of proper lipid delivery to the *C. elegans* oocyte is known to impair oocyte growth, fertilization and the development of early embryos (Kimble and Sharrock, 1983; Sturmey et al., 2009). Thus, together with yolk particles, LDs likely regulate lipid metabolism in oocytes and embryos. However, the exact contribution of LDs during oocyte and embryonic development in *C. elegans* remains largely obscure. Although the intestine is a major LD storage site in *C. elegans*, it will be of great interest to determine whether the regulation of LDs and LD-associated proteins, such as SEIPIN, share a similar genetic pathway in non-adipose tissues, such as the germline and young embryos.

In *C. elegans*, a multilayered extracellular matrix, the eggshell, begins to form when an oocyte enters the spermatheca where it is fertilized. The synthesis of this eggshell occurs in a hierarchical pattern, with each layer being sequentially added, starting with the outermost layer first. A lipid-rich layer known as the permeability barrier is the fifth layer of the eggshell to be synthesized and is responsible for preventing osmotic and mechanical stress from harming developing embryos (Johnston and Dennis, 2012; Olson et al., 2012; Stein and Golden, 2018). Embryos lacking this layer are osmotically sensitive. The inhibition of a large number of genes, either by mutation or RNAi, cause the Osmotic Integrity Defective (OID) phenotype. Many of these genes are involved in lipid biosynthesis (Benenati et al., 2009; Rappleye et al., 2003; Tagawa et al., 2001). Interestingly, depletion of the yolk receptor protein RME-2 causes the OID phenotype, suggesting that yolk lipids may contribute to the permeability barrier (Gunsalus et al., 2005). Although the specific lipids and protein components of the permeability barrier remain unknown, these earlier studies would suggest that lipid bodies, such as LDs and yolk, may contribute to permeability barrier formation. Determining the mechanisms by which the lipids get to, and help form, the permeability barrier will certainly shed light on the additional mechanisms that may operate to support fat turnover and lipid synthesis during the formation of this extracellular lipid-rich layer.

Here, we made the observation that loss of SEIP-1 in *C. elegans* not only models BSCL2 symptoms with marked lipodystrophy but also exhibit severe defects in the formation of the lipid-rich permeability barrier of the eggshell. Homozygous *seip-1* mutants produce osmotically sensitive embryos, many of which fail to hatch. Taking advantage of high resolution confocal and transmission electron microscopy, our data indicate that depletion of SEIP-1 causes abnormally sized LDs in oocytes and embryos as well as a defective permeability barrier. However, SEIP-1 depletion did not affect yolk particle synthesis, suggesting SEIP-1 may independently regulate LDs during permeability barrier formation. PUFA biosynthetic genes were also transcriptionally misregulated in our *seip-1* mutants, suggesting a crucial role of SEIP-1 for maintaining FA homeostasis during embryogenesis. Importantly, supplementing the diet of these *seip-1* mutants with PUFAs resulted in the rescue of the OID phenotype, and its associated embryonic lethality. Therefore, our study provides a powerful *in vivo* system to investigate the detailed mechanism of SEIP-1 in modulating lipid metabolism and homeostasis during early embryonic development. Specifically, maturing oocytes require SEIP-1 to help in the synthesis of the extracellular permeability barrier lipids that are made shortly after oocytes are fertilized.

## Results

### *C. elegans seip-1* complements *S. cerevisiae sei1*Δ mutants

A sole *SEIPIN1* ortholog, *seip-1 (R01B10.6)*, was identified in the *C. elegans* genome. SEIP-1 shares ∼20% amino acid identity with both human SEIPIN (BSCL2) and *S. cerevisiae* SEIPIN (Sei1p). To determine if *C. elegans* SEIP-1 could complement budding yeast *sei1Δ* strain, *C. elegans* SEIP-1 fused with a GFP tag was expressed in yeast *sei1Δ* mutants. Depletion of Sei1 protein in yeast cells caused an alteration in lipid droplet morphology, including enlarged LDs (Fig. S1A-D, M) and small clustered LDs (Fig. S1M). Expression of GFP-Seip-1p was found to either surround the LDs labeled by Erg6p-mCherry or localize adjacent to the LDs (Fig. S1E-H). Co-localization of GFP-Seip-1p with ER marker Sec63p-mCherry was observed in yeast cells (Fig. S1I-L), which is consistent with the observation that SEIP-1::mScarlet expresses at ER-LD contacts in *C. elegans* embryos (Fig. 1A) (Cao et al., 2019). Interestingly, GFP-Seip-1p significantly rescued the aberrant LD population in the *sei1Δ* yeast cells (Fig. S1E-H, M), suggesting expression of the heterologous *C. elegans seip-1* gene functionally complements the loss of *SEI1* gene in yeast cells.

**Figure 1.**
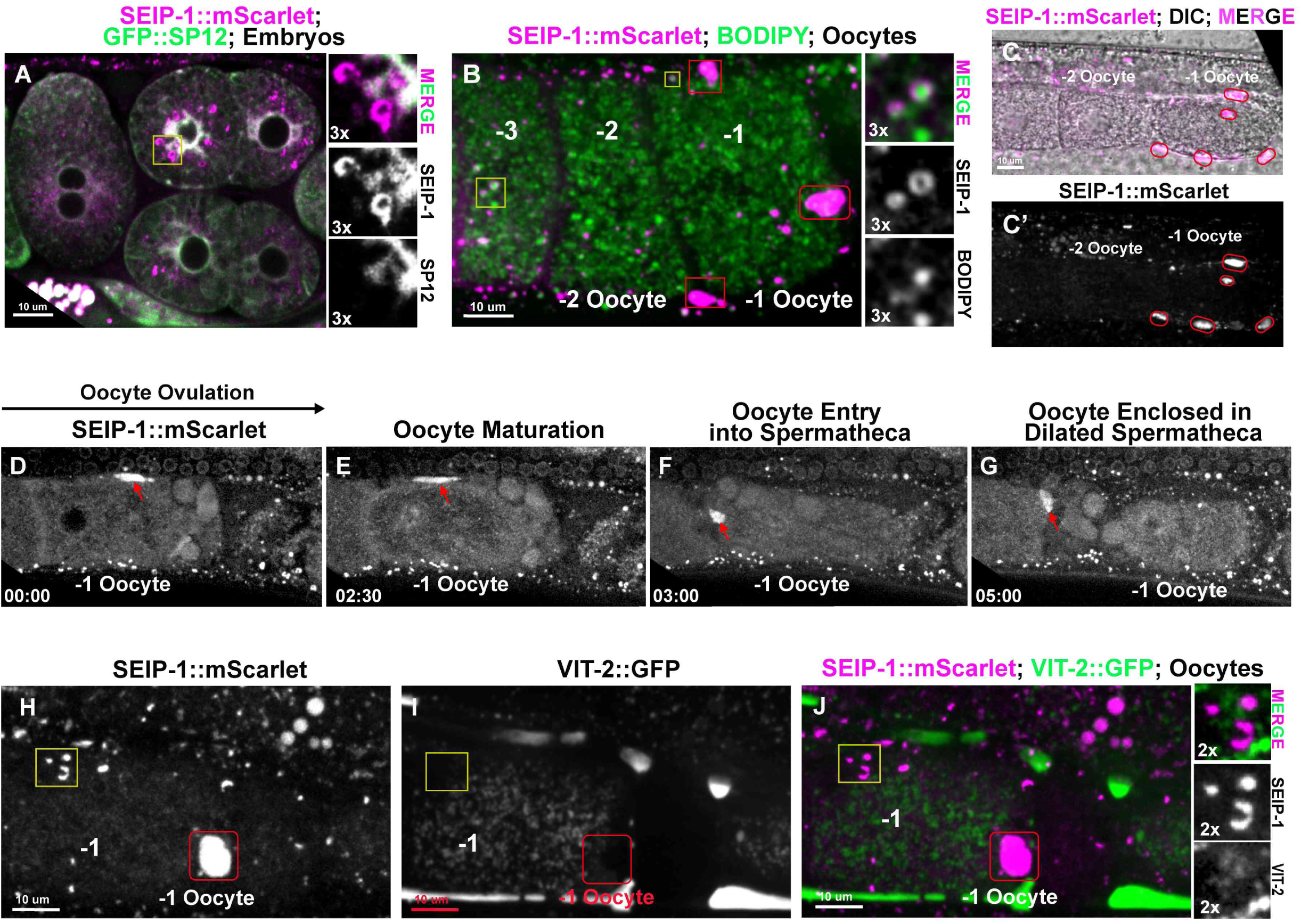
*seip-1* expression pattern in *C. elegans* oocytes and embryos. *seip-1* is expressed in *C. elegans* reproductive germline cells and embryos. (A) mScarlet was knocked-in to the *seip-1* C-terminus. SEIP-1::mScarlet is strongly expressed in the embryos, especially at ER region (yellow square). Right inserts show SEIP-1::mScarlet (magenta, middle panel) labeled circles formed at the region adjacent to the ER, labeled by GFP::SP12 (green, bottom panel). The merge image is shown in the top panel. (B) In the oocytes, SEIP-1::mScarlet labeled circular organelles surround the lipid droplets stained by BODIPY. −1 oocyte represents the oocyte most proximal to the spermatheca, while −3 oocyte represents the oocyte more distal to the spermatheca. Right inserts show SEIP-1::mScarlet (middle panel), BODIPY (bottom panel), and merged image (top panel) of the oocyte lipid droplets (yellow square in the −3 oocyte). (C-C’) SEIP-1::mScarlet (magenta in Fig. 1C) strongly expresses in the pseudocoelomic space (red squares in Fig. 1B; red circles in Fig. 1C-C’) surrounding the ovulating oocytes (−1 oocyte). (D-G) Representative images of SEIP-1::mScarlet localization during ovulation and fertilization. SEIP-1::mScarlet strongly expresses in the fifth sheath cell (red arrows) that surrounds the −1 oocyte (D) until the oocyte is ovulated and enters the spermatheca (E-G). (H-J) A vitellogenin::GFP fluorescent protein (VIT-2::GFP, green) transgene was used to monitor yolk lipoprotein localization. There is no co-localization of SEIP-1::mScarlet (H) and VIT-2::GFP (I) either in the lipid droplet (yellow square) or the pseudocoelomic space (red square). (J) Amplified images showing a lack of co-localization of SEIP-1::mScarlet (middle panel, right) and VIT-2::GFP (bottom panel, right) at the lipid droplets. The merge image was shown at the top right panel. Scale bars are indicated in each panel.

### SEIP-1::mScarlet expresses in multiple *C. elegans* tissues and on lipid droplets

To investigate the roles that SEIP-1 may play during *C. elegans* development, we investigated its expression pattern. High expression of *seip-1* mRNA was detected in the gonad, intestine, and embryos of wildtype animals (Fig. S2A). To visualize the expression pattern of SEIP-1 in *C. elegans*, we directly knocked-in a monomeric red fluorescent report gene, mScarlet, into the C-terminus of the *seip-1* endogenous locus using CRISPR/Cas9. The C-terminal fusion protein, SEIP-1::mScarlet, was widely expressed from embryonic stages through adulthood (Fig. 1A-H, J and Fig. S2B-J). This is consistent with the expression pattern of *seip-1* mRNA in these tissues (Fig. S2A). Animals with C-terminally tagged SEIP-1 did not show any embryonic lethality nor any obvious behavioral defects, suggesting that tagging SEIP-1 at the C-terminus does not disrupt function. We also attempted to knock-in both GFP and mScarlet at the N-terminus of SEIP-1, however, N-terminal insertions caused embryonic lethality and no expression of fluorescent proteins. These observations suggest that N-terminal insertions disrupt SEIP-1 functions. Since SEIPIN1 is an integral endoplasmic reticulum (ER) membrane protein, an ER marker, GFP::SP12 (Poteryaev et al., 2005), was expressed in the SEIP-1::mScarlet animals to determine whether SEIP-1 associates with ER subdomains. Indeed, SEIP-1::mScarlet and GFP::SP12 clearly co-localize in the regions surrounding embryonic nuclear envelopes (Fig. 1A). Circular structures of SEIP-1::mScarlet formed near the ER, consistent with the idea that SEIPIN resides in the ER membrane to regulate nascent LD biogenesis (Fig. 1A). Since SEIPIN1 has been proposed to promote the maturation of nascent LDs, we stained animals expressing SEIP-1::mScarlet with the lipophilic dye BODIPY 493/503, which stains LDs. We observed in maturing oocytes that SEIP-1::mScarlet was present on ring structures surrounding a subset of BODIPY-stained LDs (Fig.1B and inserts). Similar co-localizations were found in the embryos and different tissues of larva and adult animals (Fig. S2C-G). Strong expression of SEIP-1::mScarlet was observed in the fifth gonadal sheath cell that surrounds the maturing oocytes prior to ovulation into the spermatheca (Fig. 1B-H, J, Video S1). In *C. elegans*, the pseudocoelomic lipoprotein pool was identified as yolk, which is secreted from the intestine and travels through the gonadal basal lamina and sheath pores and then is endocytosed by the developing oocytes (Paupard et al., 2001). To test whether the expression of SEIP-1::mScarlet regulates yolk delivery to maturing oocytes, the yolk marker VIT-2::GFP was co-expressed in the animals containing SEIP-1::mScarlet (Fig. 1H-J). VIT-2::GFP was not present on the SEIP-1::mScarlet labeled droplets in the intestine and oocytes (Fig. 1H-J). Taken together, SEIP-1::mScarlet accumulated at neutral lipid-containing droplets, which were not characterized by the presence of the yolk marker. SEIP-1::mScarlet strongly expressed in the oocytes and sperm (Fig. S2H-I), which suggested that SEIP-1 may play roles in *C. elegans* reproduction. To our knowledge, this is the first report of SEIPIN in both gametes and embryos of *C. elegans* and may represent a new *in vivo* model to study SEIPIN function in non-adipose tissues, particularly in germ cells and reproductive tissues.

### *seip-1* mutants disrupt permeability barrier formation in newly fertilized embryos

To investigate the biological functions of SEIP-1 in *C. elegans*, three candidate null deletion alleles were generated by CRISPR/Cas9 genome editing; one allele was a deletion of the whole *seip-1* gene [*seip-1(av109)*] (Fig. 2A, Fig. S3C, G) and the other two alleles [*seip-1(cwc1)* and *seip-1(cwc2)*] were generated by replacing the *seip-1* gene with an expression cassette of a hygromycin B resistance gene (*HygR*) (Radman et al., 2013) (Fig. 2B, Fig. S3B, D-F). A small *seip-1* deletion allele, *seip-1(tm4221)*, was also tested in this study (Fig. S4C). The number of F1 progeny (brood size) was significantly reduced in all tested homozygous deletion mutants compared with wild type (Fig. 2C). In addition, the embryonic viability of F1 progeny was dramatically decreased in all tested *seip-1* mutants compared with wild type. On average, 10%-35% of the tested *seip-1* mutant embryos failed to hatch (Fig. 2D). Expression of a *seip-1* cDNA plasmid in the *seip-1* mutants significantly rescued the severe embryonic lethality of F1 progeny, suggesting the embryonic lethality was due to the deletion of *seip-1* (Fig. S3H). Surprisingly, we did not observe any obvious defects during later developmental processes in the hatched *seip-1* mutant larvae (data not shown), suggesting *seip-1* may only have an essential function during early embryogenesis. It has been suggested that *de novo* lipid machinery is critical for early embryonic development while later larval development utilize fatty acid precursors from their bacterial diet (Watts et al., 2018).

**Figure 2.**
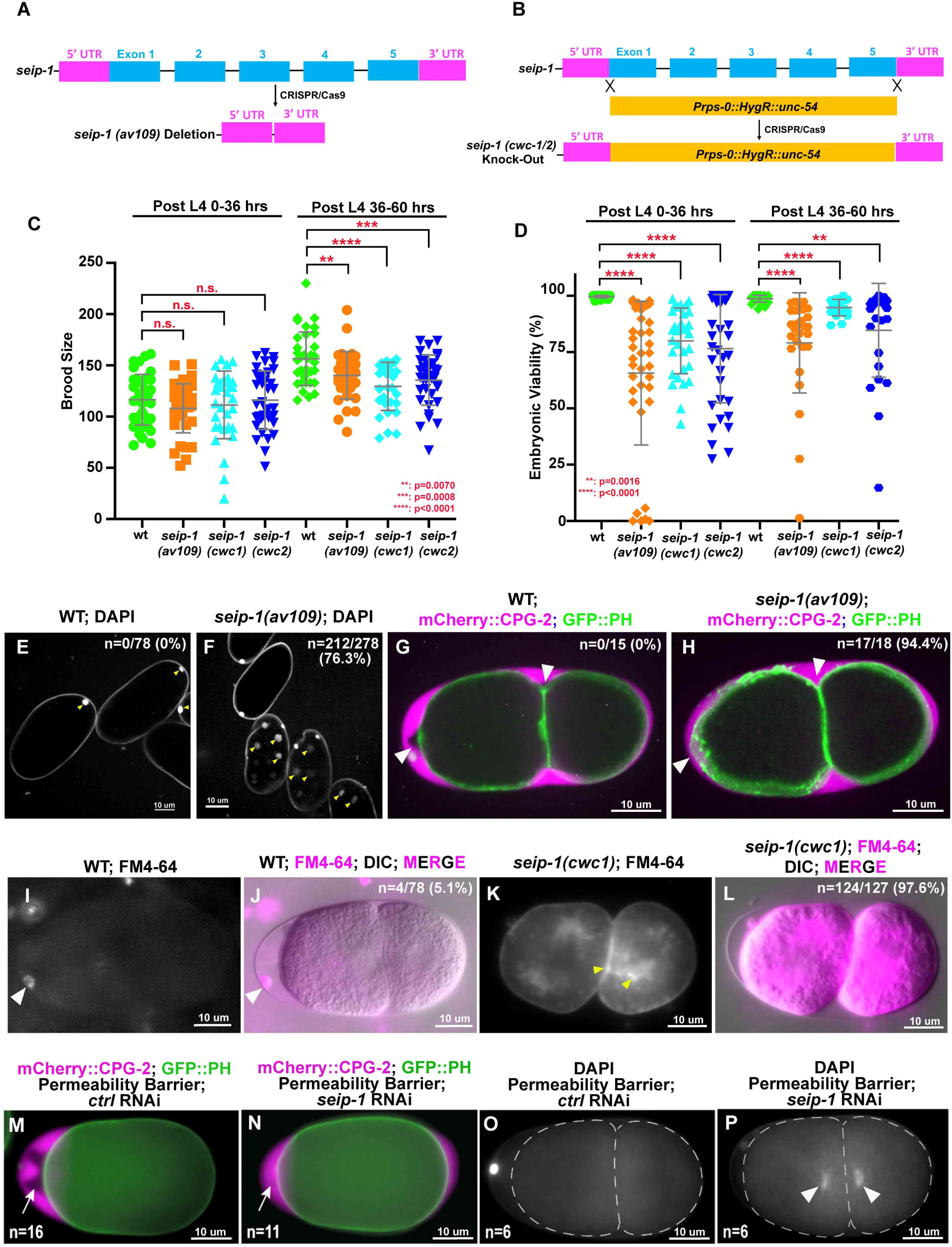
Deletion of the *seip-1* gene causes severe eggshell formation defects. (A) Diagram of deleted regions in *seip-1(av109)* mutant or (B) replacement of *seip-1* gene with a hygromycin-resistant gene (*HygR*) cassette in *seip-1(cwc1)* and *seip-1(cwc2)* mutants. (C) Brood size was significantly reduced in *seip-1* mutants after 36 hours post mid-L4 (Day 2-3 adults). (D) The percentage of viable embryos was reduced in the *seip-1* deletion animals. (E, F) DAPI staining of zygotic chromatin (yellow arrows, F) was apparent in *seip-1* embryos, while only first polar body (yellow arrows, E) stained in wildtype embryos (n=number of embryos with zygotic chromatin stained with DAPI / number of embryos imaged). (G, H). mCherry::CPG-2 (magenta) was used to visualize the integrality of the permeability barrier. In wild type, the inner layer of the eggshell prevents diffusion of mCherry::CPG-2 beyond the space between outer layers of the eggshell and the embryo surface (white arrows, G). However, in *seip-1(av109)* embryos, mCherry::CPG-2 penetrates inside the entire space between eggshell and the embryo surface (H; n=number of embryos with mCherry::CPG-2 penetration / number of embryos imaged). (I, J) Only the plasma membrane surrounding the first polar body is stained by FM4-64 (magenta) in wildtype (WT) embryos, while staining was observed at the plasma membrane and cytosol in *seip-1(cwc1)* mutant embryos (K, L) (n=number of embryos with FM4-64 penetration / number of embryos imaged). (M, N) Depletion of SEIP-1 by RNAi displayed defective permeability barrier formation, allowing mCherry::CPG-2 (white arrows, M, N) to fill the space between the outer layers of the eggshell and the embryo surface (n=number of embryos with mCherry::CPG-2 penetration). (O, P) DAPI (white arrowheads, P) penetrates inside the embryos to stain zygotic chromatin (n=number of embryos with DAPI penetration). Scale bars are indicated in each panel. P-values: ** = 0.0070 (C); ** = 0.0016 (D); *** = 0.0008 (C); **** <0.0001 (C, D) (t-test).

Many of the *seip-1* mutant embryos displayed a shrunken and misshapen morphology, which is suggestive of eggshell formation defects (Fig. S4A, B). Since small molecules, such as DAPI and the lipophilic dye FM4-64, are able to penetrate into fertilized embryos with defective eggshells, *seip-1* mutant embryos were incubated with DAPI and FM4-64 to test the integrity of their eggshells. Zygotic chromatin, labeled by DAPI, and cytoplasmic membranes, labeled by FM4-64, were frequently observed in the post-meiotic embryos of the *seip-1* mutants (Fig. 2F; K, L). In contrast, only the first polar body was stained by DAPI and no membranes were stained by FM4-64 in wildtype embryos (Fig. 2E, I, J). Additionally, mCherry::CPG-2 was used to visualize the permeability barrier in this study. In wild type, the lipid-rich permeability barrier, which defines the innermost layer of the eggshell, prevents diffusion of mCherry::CPG-2 beyond the space between the outer layers of the eggshell and the permeability barrier (Fig. 2G) (Olson et al., 2012). The white arrowheads highlight the edge of the permeability barrier that normally excludes mCherry::CPG-2. However, in *seip-1* null mutants, the permeability barrier is disrupted and mCherry::CPG-2 fills the entire space between the outer eggshell and the embryonic surface (Fig. 2H). The depletion of SEIP-1 by RNAi and a small deletion *seip-1(tm4221)* mutant displayed identical eggshell permeability defects (Fig. 2M-P, Fig. S4D, E). Collectively, these data suggested that loss of SEIP-1 caused severe defects in eggshell formation, which may account for the embryonic lethality observed in these mutants.

To characterize the eggshell defects of the *seip-1* mutants in greater detail, we used thin-sectioned electron microscopy (EM) to examine the ultrastructure of forming eggshells in embryos, which reside in the uteri of gravid hermaphrodites. Previous studies demonstrated that eggshell formation occurs in a hierarchical pattern, with each layer being sequentially added (Olson et al., 2012) starting from the outermost layer. The whole process starts from the unfertilized oocyte (which is at prophase of meiosis I) and ends at anaphase II of the meiotic embryo (Fig. 3A). A lipid-rich layer known as the permeability barrier is the fifth layer of the eggshell, which is formed during anaphase of meiosis II (Johnston and Dennis, 2012; Olson et al., 2012; Stein and Golden, 2018) (Fig. 3A). The examined embryos were labeled as +1 to +5, which represents the positioning from the distal end of uterus (+1; adjacent to the spermatheca) to the vulva (+5) (Fig. 3A). EM images indicated that the youngest newly-fertilized +1 embryo has an outermost vitelline layer (synthesized pre-fertilization) and a chitin layer (formed right after fertilization but before anaphase I of meiosis), while the +2 embryo shows an additional forming chondroitin proteoglycan (CPG) layer (fully formed after anaphase of meiosis I), in both wildtype and *seip-1(cwc1/2)* mutants (Fig. 3B, C, E-G). The permeability barrier was initially synthesized in the next oldest embryos (during anaphase of meiosis II) and properly formed in the older embryos (approximately post anaphase II of meiosis) in wildtype animals (Fig. 3D, G). All of the examined mitotic embryos (+3 to ∼ +5 embryos) contain a properly formed permeability barrier (Fig. 3G) in wildtype embryos. However, we could not identify the permeability barrier structure in any *seip-1(cwc1/2)* embryos, including in the mitotic embryos (Fig. 3F, G). Only the first three outer layers, vitelline layer (VL), chitin layer (CL) and CPG layer (CPGL) were observed in *seip-1* mutant embryos. This is consistent with the observation that the vitelline layer marker, PERM-4::mCherry (Gonzalez et al., 2018), and CPG layer marker, mCherry::CPG-1, are properly localized when *seip-1* was depleted by RNAi (Fig. 3H-K). In sum, our data strongly support the conclusion that SEIP-1 is essential for the proper formation of the permeability barrier layer of the eggshell.

**Figure 3.**
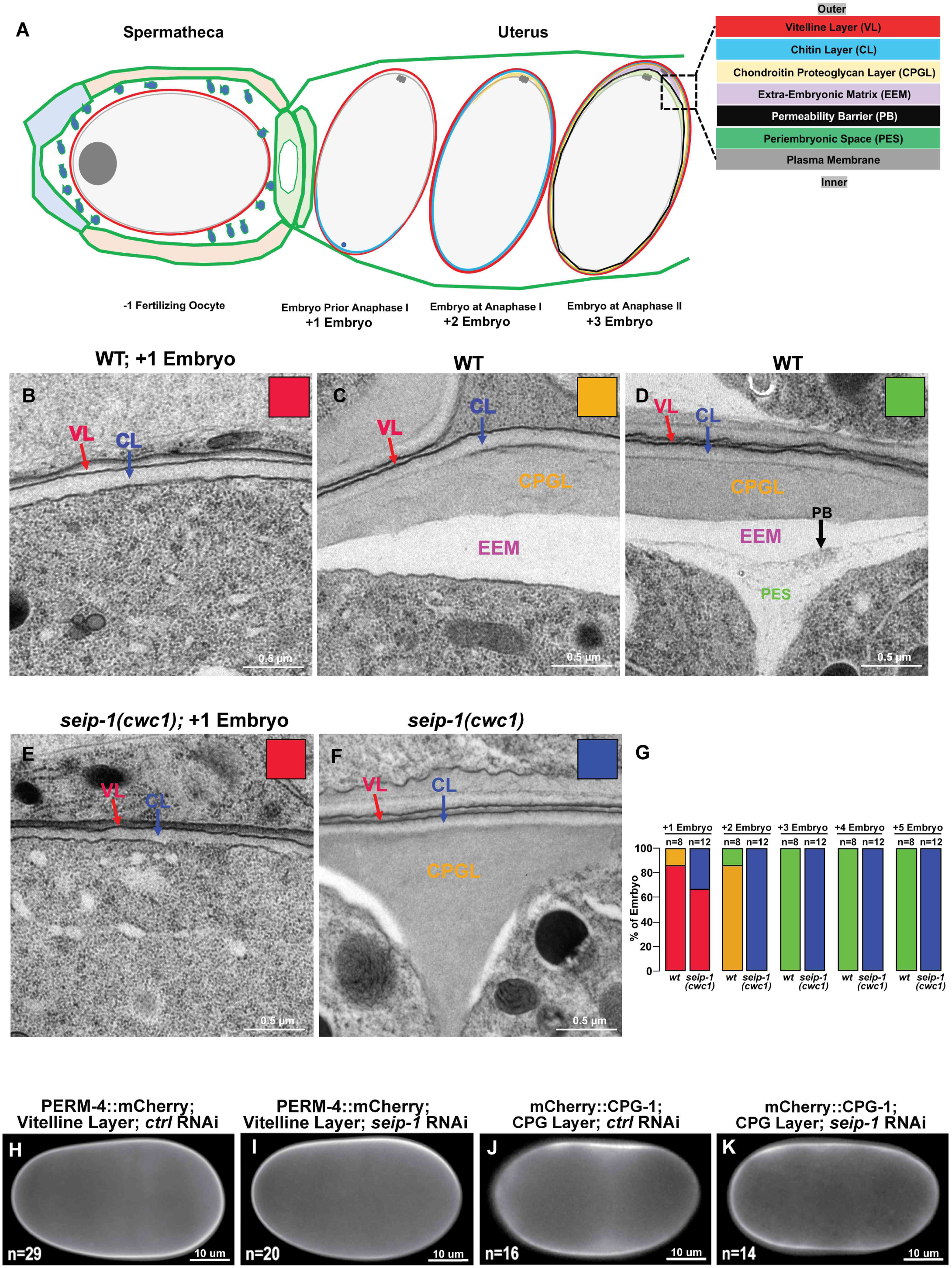
Formation of the permeability barrier requires functional SEIP-1. (A) Diagram of sequential steps in assembly of the eggshell. At anaphase of meiosis II, the permeability barrier is formed between the outer eggshell and the plasma membrane. (B, C, E) Transmission electron micrographs of high-pressure frozen embryos demonstrate that the vitelline and chitin layers are easily observed in both WT and *seip-1(cwc1)* embryos that have completed meiosis I. (D) A clear edge of the permeability barrier (PB, black arrow) was visualized in every WT embryo that completed meiosis II. However, no permeability barrier was found in any stage *seip-1(cwc1)* embryos (E, F). (G) Quantification of the embryos with the forming eggshell in both WT and *seip-1(cwc1)* embryos. The embryos containing different components of the eggshell were categorized into four groups, which are represented by the colored squares inserted at the top right of each panel (B-F). Red square represents the embryos containing vitelline and chitin layers; blue squares represent the embryos with vitelline, chitin and chondroitin proteoglycan layers (CPGL); orange squares represent the embryos with four layers, including VL, CL, CPGL and extra-embryonic matrix (EEM); and the green squares represent the embryos with all five eggshell layers (VL, CL, CPGL, EEM, and PB). (H-K) At the light microscopy level, the eggshell proper appears mainly intact when SEIP-1 was depleted by RNAi, as markers for the vitelline layer (PERM-4::mCherry, H, I) and CPG layer (mCherry::CPG-1, J, K) are properly localized compared with embryos treated by control RNAi (n= the number of embryos examined). Scale bars are indicated in each panel.

### *seip-1* mutants have abnormal lipid droplets

Previous studies identified the eggshell’s permeability barrier as a lipid-rich layer, in which glycolipids may play an important role in its formation (Olson et al., 2012; Stein and Golden, 2018; Watts et al., 2018). However, the delivery and secretion mechanisms of the lipids required to build the permeability barrier have not been characterized to date. To investigate whether the defective formation of the eggshell in *seip-1* mutants was caused by aberrant LDs, we performed vital BODIPY staining to mark the lipid droplets that surround maturing oocytes and nascent embryos. All three *seip-1* deletion alleles showed enlarged LDs in the oocytes and embryos (Fig. 4A-D, S5A). Greater than 10 enlarged LDs (>1.5 µm) were observed in the −1 to −3 oocytes in each of these mutants (Fig. 4D, S5A). In contrast, there were no enlarged LDs observed in oocytes in wild type (Fig. 4D, S5A). Additionally, EM analysis showed that the abnormal LDs exist in both *seip-1(cwc1)* oocytes and embryos (Fig. 4E-H, E’-H’). Consistent with a previous study (Cao et al., 2019), a large number of tiny LDs and enlarged LDs were also observed in the major fat storage tissue, the intestine, in the *seip-1* deletion mutants (Fig. S5B). Overall, these observations are consistent with previous studies in mammalian and yeast cells that dysfunction of *seip-1* causes aberrant LD morphology, including both enlarged and abnormally small LDs (Fei et al., 2008; Grippa et al., 2015; Szymanski et al., 2007; Wang et al., 2016a; Wolinski et al., 2011).

**Figure 4.**
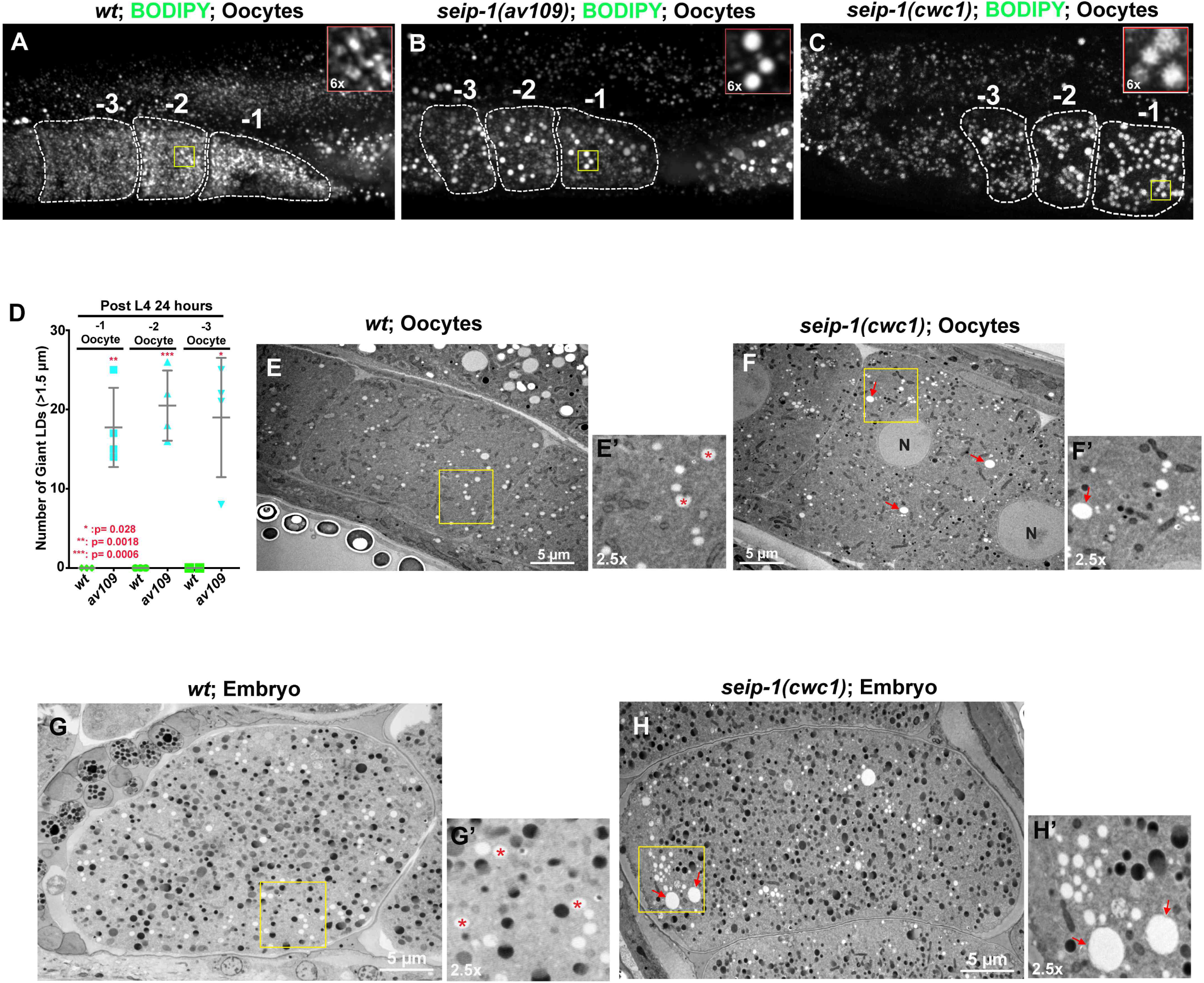
SEIP-1 depletion alters lipid droplet (LD) size in the oocytes and embryos. (A-C) BODIPY-stained LDs in the −1 to −3 oocytes of wildtype and *seip-1* animals. Top right inserts highlight the region marked by yellow squares in each panel; these are individual LDs. (B, C) Abnormally enlarged LDs were frequently observed in the −1 to −3 oocytes of *seip-1* mutants. (D) Quantification of the enlarged LDs (diameter > 1.5 µm) inside the −1 to −3 oocytes of both wildtype and *seip-1* mutants. (E, F) Transmission electron micrographs of high-pressure frozen oocytes show normal LDs (red asterisks in E’) in wildtype oocytes (E) and enlarged LDs (red arrows) in the *seip-1(cwc1)* oocytes (F, F’). (E’, F’) are 2.5X magnification of the regions marked with a yellow square in panels E and F showing individual LDs. (G, H) Transmission electron micrographs of high-pressure frozen embryos show normal LDs (red asterisks in G’) in wildtype embryos (G) and enlarged LDs (red arrows) in the *seip-1(cwc1)* embryos (H, H’). (G’, H’) are 2.5X magnification of the regions marked with a yellow square in panels G and H showing individual LDs. Scale bars are indicated in each panel. P-values: * = 0.028; ** = 0.0018; *** = 0.0006 (t-test).

### The fatty acid synthesis pathway is disrupted in *seip-1* mutants

The assembly of long chain fatty acids (LCFAs) requires at least seven key enzymatic reactions (Fig. 5A). To test the effect of *seip-1* deletions on the transcriptional level of each of these enzymes, quantitative PCR analysis was carried out in both wildtype and *seip-1* mutant embryos (Fig. 5B). Detailed analysis revealed that the expressional levels of FA metabolism pathway genes were affected by the dysfunction of *seip-1* in *C. elegans* embryos. These results indicated that four FA synthetic genes (*elo-2*, *fat-2*, *fat-5*, and *fat-6*) were up-regulated, while another four FA synthetic genes (*pod-2*, *elo-1*, *fat-1* and *fat-4*) were down-regulated in *seip-1* mutants compared with wild type (Fig. 5A, B). A wide range of PUFAs are synthesized in *C. elegans* through *de novo* fatty acid metabolism (Watts and Ristow, 2017). Four types of fatty acid desaturases, including Δ5 desaturase (*fat-4)*, Δ6 desaturase (*fat-3),* three Δ9 desaturases (*fat-5-7),* and a Δ12 desaturase (*fat-2)*, and two elongases, *elo-1* and *-2*, are responsible for making a range of C18 or C20 PUFAs (Fig. 5A). Quantitative PCR results further indicated that the transcriptional level of a few PUFA synthesis related enzymes, such as *elo-2*, *fat-2* and *fat-6*, were significantly up-regulated in *seip-1* deletion mutant embryos (Fig. 5A, B). In contrast, the expression of other enzymes, such as *elo-1* and *fat-4*, were reduced in the *seip-1* mutants. Interestingly, the genes *acox-1.1*, *acox-3*, *daf-22*, and *dhs-28,* which play roles in fatty acid β-oxidation to break down fatty acids to generate acetyl-CoA, were significantly up-regulated in *seip-1* mutants (Fig. 5B).

**Figure 5.**
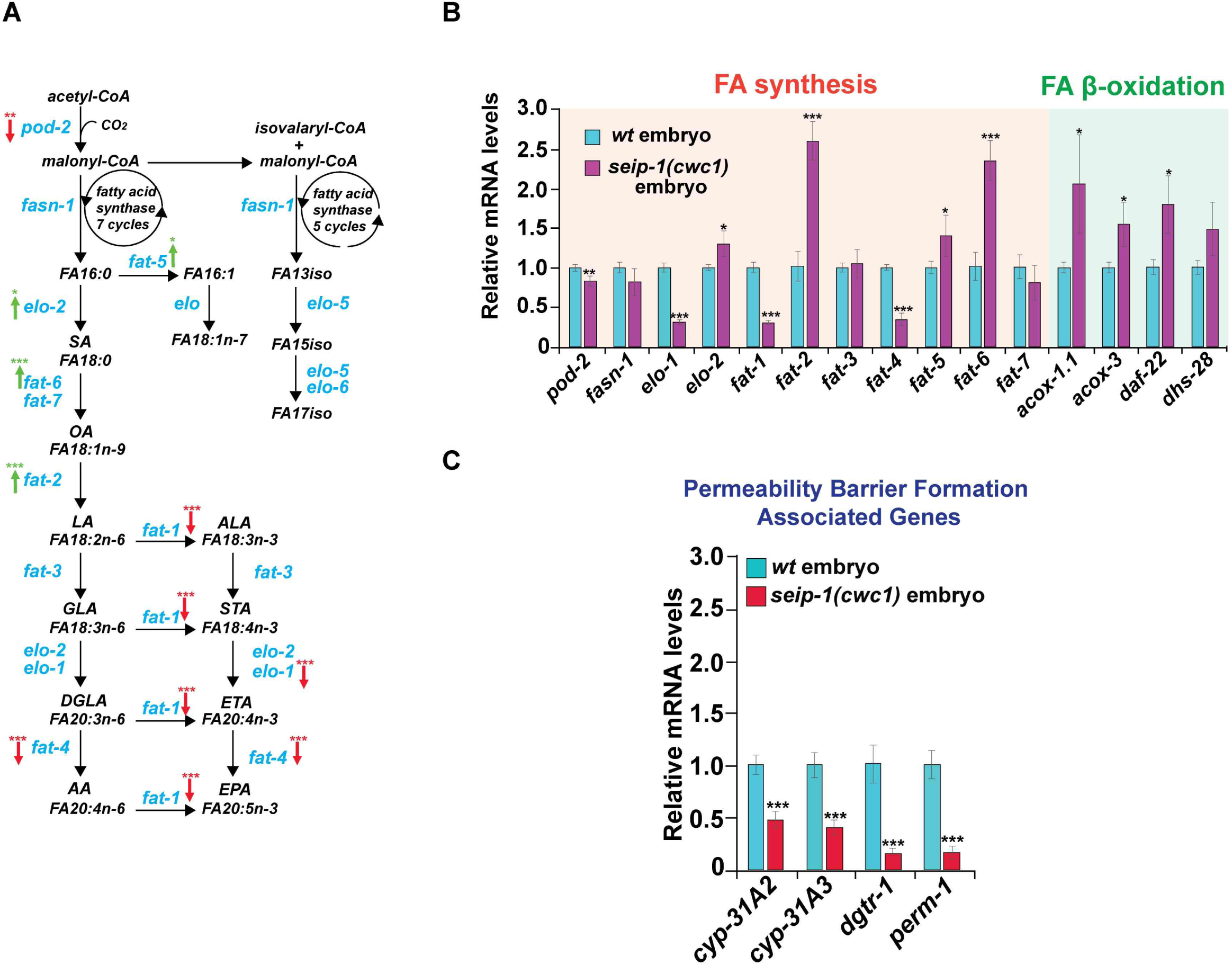
Gene expression of some fatty acid synthesis genes are mis-regulated in *seip-1* embryos. (A, B) Expression of the *de novo* fatty acid synthetic genes were either up-regulated (*elo-2*, *fat-2*, *fat-5* and *fat-6*, red arrows in A) or down-regulated (*pod-2*, *elo-1*, *fat-1*, *fat-4*, green arrows) in *seip-1(cwc1)* mutant embryos. FA β-oxidation gene expression was up-regulated in *seip-1(cwc1)* mutant embryos (B). Quantitative transcriptional expression of these genes was determined by qRT-PCR. (C) Genes associated with the formation of the permeability barrier were down-regulated in *seip-1* mutant embryos compare with wildtype embryos. P-values: * < 0.05; ** < 0.01; *** = 0.001 (t-test).

To examine the global effect on gene expression, cDNA microarray analysis was carried out in both wildtype and *seip-1* mutant embryos (Fig. S6A). Testing 20,221 genes with cDNA microarray analysis revealed the global transcriptional variations in *seip-1* mutant embryos compared with wildtype embryos (Fig. S6A). 8871 of 20,221 genes displayed either up-regulated or down-regulated changes by at least 1.5-fold with P value <0.05 in *seip-1* mutant embryos. In contrast, only a small population of genes (622 of 20,221) was affected by loss of SEIP-1 in whole animals. Additionally, Kyoto Encyclopedia of Genes and Genomes (KEGG; www.genome.jp/kegg/) (Kanehisa et al., 2019) pathway enrichment analyses identified metabolic pathways, including FA metabolism pathways, were being affected the most in the *seip-1* mutants embryos compared with wild type (Fig. S6B). Taken together, loss of SEIP-1 may disrupt *de novo* FA synthesis and FA homeostasis during embryogenesis. Future studies will decipher the precise mechanism of how SEIP-1 influences fatty acid homeostasis.

The *C. elegans* permeability barrier was identified as a lipid-rich layer, which primarily consists of fatty acid derivatives (Olson et al., 2012), and the current study links lipid droplets to permeability barrier formation. Triacylglycerol (TAG) and sterol esters (SE) are commonly stored in LDs, and genes implicated in TAG and SE synthesis have been shown to give rise to the Osmotic Integrity Defective (OID) phenotype and impaired permeability barrier formation when depleted by RNAi (Olson et al., 2012). The DGAT2-related acyltransferase DGTR-1 may convert diacylglycerol (DAG) to TAG, while the cytochrome P450s CYP-31A2/A3 and the putative hydroxysteroid dehydrogenase/isomerase PERM-1 enzyme could theoretically modify sterol esters. In *seip-1* mutant embryos, transcriptional expression of *cryp-31A2/3*, *perm-1* and *dgtr-1*, were tested by qRT-PCR and found to be significantly down-regulated compared with wildtype embryos, suggesting that dysfunction of *seip-1* may disrupt synthesis of lipids traditionally stored within lipid droplets (Fig. 5C). This defective lipid synthesis is likely due to the abnormal or unbalanced fatty acid content observed in *seip-1* mutants. Therefore, the defective permeability barrier formation in *seip-1* mutants is likely associated with a deficiency of lipids stored within lipid droplets.

### Dietary supplementation of PUFAs rescued the permeability barrier defects

From our gene expression studies, we hypothesized that diminished PUFA synthesis could, in part, be responsible for the defective formation of the permeability barrier in *seip-1* mutants. *fat-*2 embryos have similar permeability defects and have been previously rescued by PUFA supplementation (Watts et al., 2018). To address this hypothesis, different PUFAs were added to the diet of the *seip-1* mutants. Feeding animals 300 µM ω-6 PUFA, dihomo-gamma-linolenic acid DGLA (C20:3_n-6_), significantly rescued the embryonic lethality in all three *seip-1* deletion mutants (Fig. 6A). Although brood sizes in the tested animals were reduced in the PUFA-fed animals (data not shown), greater than 95% of these mutant embryos hatched after PUFA dietary supplementation. To determine whether the defective permeability barrier was also restored, different reporter genes were assessed in the PUFA-fed *seip-1* mutant animals. In the mutants alone, defective permeability barrier formation led to mCherry::CPG-2 filling the entire space between the outer eggshell and the embryonic surface (Fig. 6B, C). Small molecules, such as DAPI and FM4-64, were able to penetrate inside the eggshell and stain the zygotic chromatin and plasma membranes, respectively (Fig. 6B, D, F). No obvious permeability barriers were observed under DIC microscopy in the mutants (Fig. 6E). However, with dietary supplementation of PUFAs, a significantly higher percentage of *seip-1* mutant embryos formed proper permeability barriers to prevent the penetration of DAPI, FM4-64, and mCherry::CPG-2 into the cytosol and peri-embryonic space (Fig. 6B’-D’, F). A clear permeability barrier was also observed by DIC microscopy (Fig. 6E’). EM imaging further confirmed that the defective permeability barriers were significantly rescued in the *seip-1(cwc1)* mutant after dietary supplementation of DGLA (C20:3_n-6_) (Fig. 6G-I). Linoleic acid (C18:2_n-6_) less efficiently rescued the permeability barrier defects when fed to the *seip-1* mutants (Fig. 6F, H). Unexpectedly, the number of enlarged LDs was slightly increased in both wildtype and *seip-1(av109)* oocytes after supplementation with the PUFA DGLA (C20:3_n-6_) (Fig. 6J-L, J’, K’). Collectively, our data suggested that reduced PUFA levels compromise permeability layer formation in the *seip-1* mutants, and thus led to embryonic lethality. This lethality can be reversed when the diet is supplemented with selected PUFAs. Loss of SEIP-1 may disrupt the availability of PUFAs by either disrupting their synthesis or by interfering with their ability to directly get incorporated into the permeability barrier. The embryonic lethality of the *seip-1* mutants is likely due to the malformation of a robust permeability barrier, leading to osmotic stress during embryogenesis.

**Figure 6.**
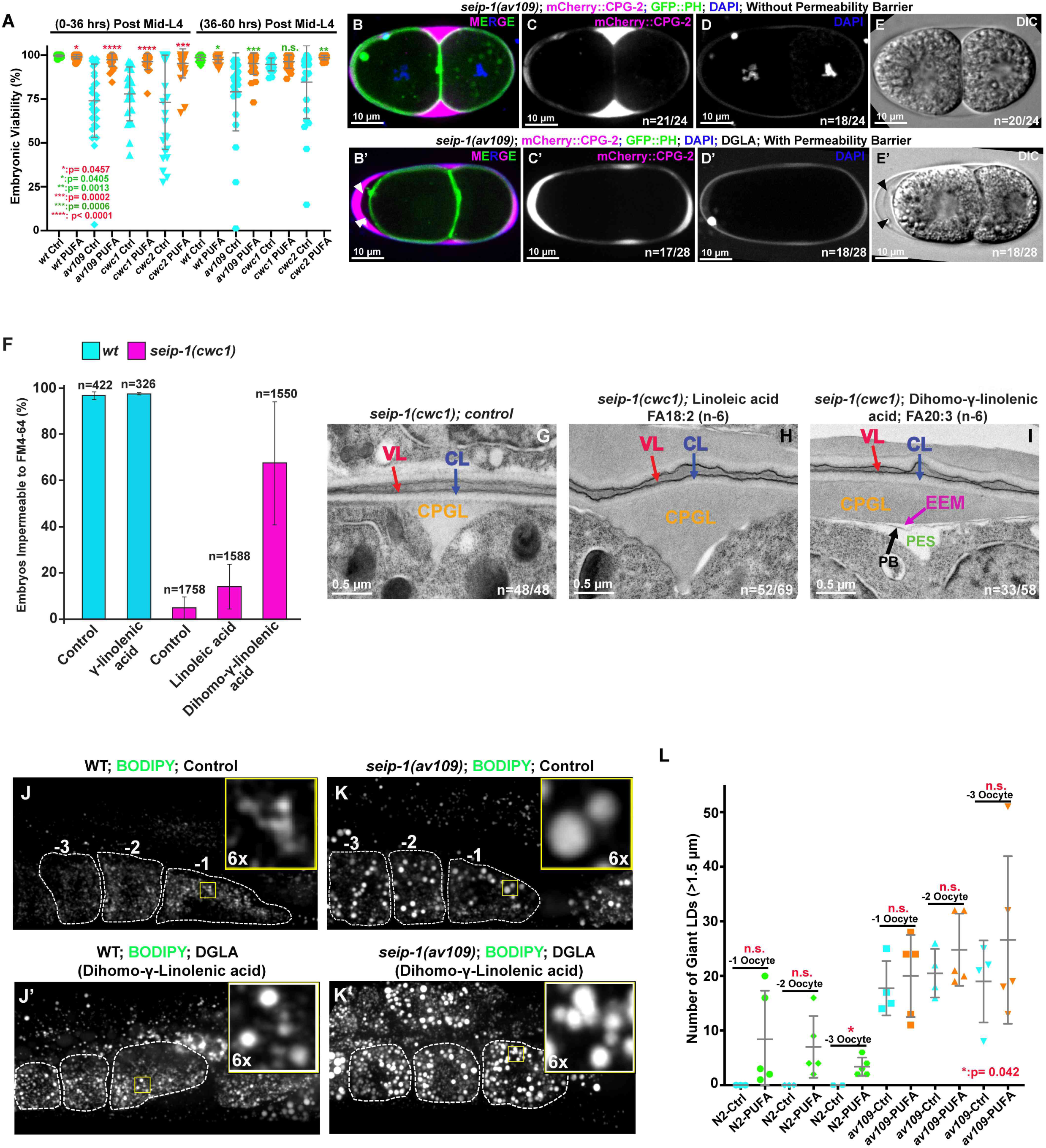
Dietary supplementation of PUFAs rescued the permeability barrier defects but not the abnormal LDs in *seip-1* mutants. (A) The percentage of viable embryos was significantly rescued in the PUFA-fed *seip-1* deletion animals (WT: green, mutant: blue, PUFA-fed: orange). (B-E) The penetration of DAPI, the expression pattern of CPG2::mCherry, and DIC images demonstrated that the permeability barrier was defective in *seip-1* mutants embryos (n=number of embryos with abnormal DAPI penetration and abnormal CPG2::mCherry pattern / number of embryos imaged) The GFP::PH transgene was used to indicate the plasma membranes. (B’-E’) The penetration of DAPI, the expression pattern of CPG2::mCherry, and DIC images demonstrated that the permeability barrier was restored after the diet of *seip-1* mutants animals was supplemented with PUFAs (n=number of embryos with normal DAPI penetration and normal CPG2::mCherry pattern / number of embryos imaged). The permeability barrier can be easily seen in panel B’ (white arrowheads) and panel E’ (black arrowheads). (F) FM4-64 staining indicated that dihomo-γ-linolenic acid performed more efficiently than linoleic acid to rescue the defective formation of the permeability barrier. (G-I) Transmission electron micrographs of high-pressure frozen embryos showing the permeability barrier (black arrow) formed after *seip-1* mutant animals were fed with dihomo-γ-linolenic acid (FA20:3_n-6_). n=number of embryos showing representative eggshell layers / number of embryos imaged. (J-K’) Abnormal LDs were frequently observed in −1 to −3 oocytes of both wildtype (J’) and *seip-1(av109)* (K’) animals fed with dihomo-γ-linolenic acid (FA20:3_n-6_). Right top inserts represent 6x amplified images of the inlaid yellow squares. (L) Quantification of enlarged LDs (diameter > 1.5 µm) in −1 to −3 oocytes of both PUFA-fed wildtype and *seip-1(av109)* animals. Scale bars are indicated in each panel. P-values: * < 0.05 (A, L); ** <0.01 (A); *** <0.001 (A); **** <0.0001 (t-test).

### Modeling human BSCL2 genetic diseases in *C. elegans*

The present study provides strong evidence that *seip-1* is essential for *C. elegans* embryogenesis, eggshell formation, and fatty acid synthesis and metabolism. Given the striking phenotypes associated with *seip-1* deletions, it is plausible to model human SEIPIN associated diseases with SEIPIN1 missense mutations using *C. elegans* embryos. Of the many missense mutations associated with disease, we chose to test a conserved alanine (p. A212P) in SEIPIN1. Patients bearing this mutation have been diagnosed with the autosomal recessive Beradinelli-Seip congenital generalized lipodystrophy type 2 (BSCL2) syndrome (Magre et al., 2001). This syndrome, like other lipodystrophies, is characterized by a total lack of adipose tissue in the body and a very muscular appearance. Fat storage thus takes place in other organs in the body, such as the liver, which can lead to hepatomegaly and liver failure (Garg, 2004; Simha and Garg, 2003). Previous studies have shown that this A212P missense mutation is a loss-of-function mutation in SEIPIN1 (Salo et al., 2016; Szymanski et al., 2007). Bioinformatics analysis indicated that alanine 185 (A185) in *C. elegans* SEIP-1 is the homologous alanine residue to human SEIPIN1 A212. Using CRISPR/Cas9, we generated this missense mutation in *C. elegans* and named the allele *seip-1(av160*[*A185P*]). We will refer to this allele as *seip-1(A185P)* for sake of clarity for the remainder of this study. To determine the phenotypic consequences of this patient-specific allele, *seip-1(A185P),* homozygous animals were examined for defects in LD size, permeability barrier defects, and embryonic lethality. Homozygous *seip-1(A185P)* displayed embryonic lethality (Fig. 7A, B), defective permeability barrier formation (Fig. 7C) and enlarged LDs (Fig. 7D, E), similar to those observed in our *seip-1* null deletion mutants. Dietary supplementation with PUFAs significantly rescued the lethality of *seip-1(A185P)* embryos laid by mothers that were fed PUFAs on day 1 (0-36 hours post L4) (Fig. 7F). However, day 2-3 (36-60 hours post L4) PUFA-fed *seip-1(A185P)* animals revealed an increase in embryonic lethality compared with wild type (Fig. 7F), suggesting that the patient-specific allele is sensitive to excessive PUFAs. Overall, these observations strongly support the idea that *C. elegans* is an appropriate model system to study SEIPIN-associated diseases. We will test other patient-specific alleles that alter residues that are conserved in the *C. elegans* SEIP-1 protein. Using such alleles, we will perform suppressor screens to help identify other potential genetic interactors that restore viability to *seip-1* embryos.

**Figure 7.**
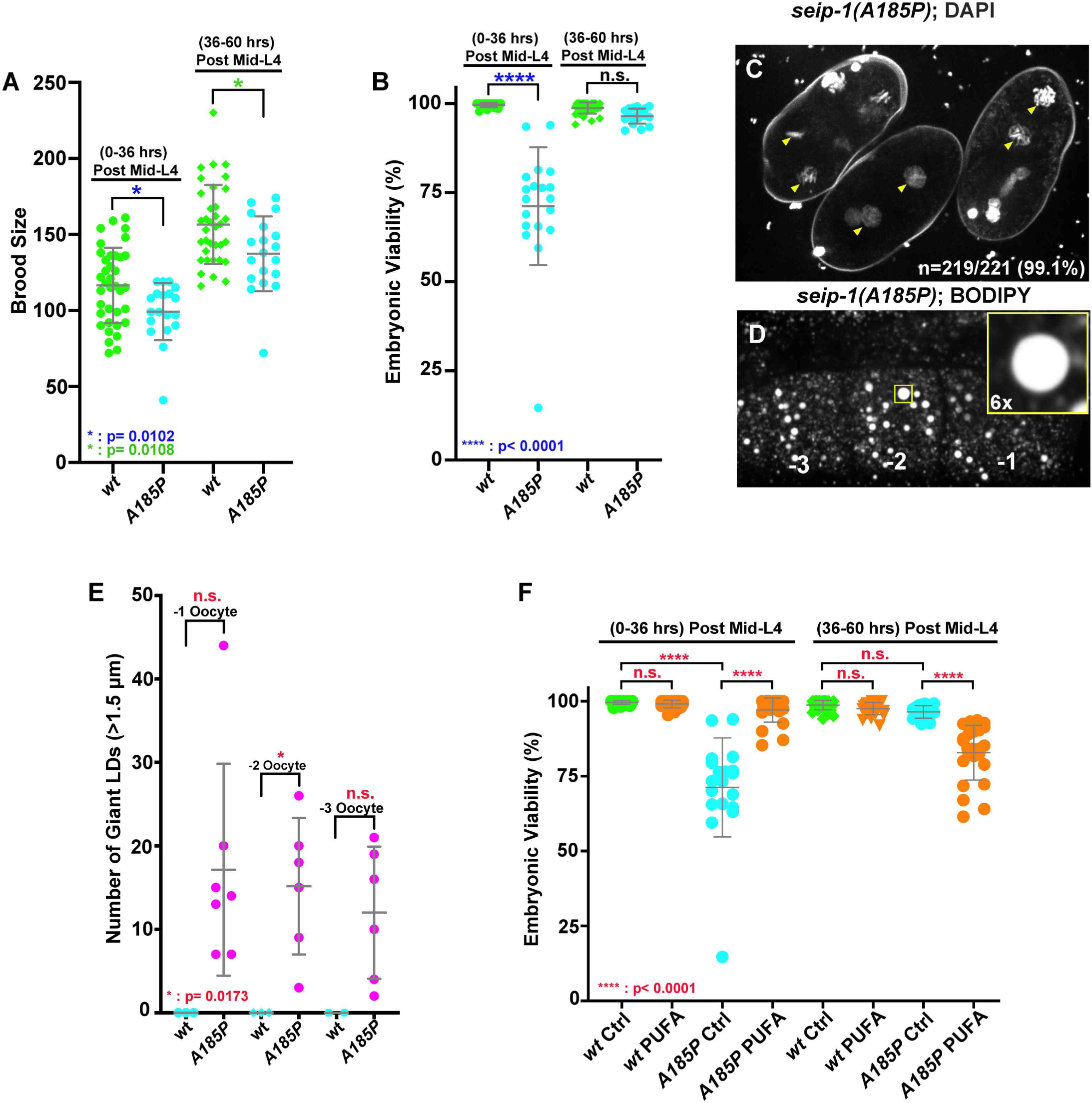
A *SEIPIN* patient-specific allele disrupts eggshell formation and displays enlarged LDs in *C. elegans*. (A) A conserved patient-specific allele *seip-1(A185P)* was generated and results in reduced brood sizes. (B) *seip-1(A185P)* causes severe embryonic lethality in the day 1 (post L4 0-36 hours) adult animals. However, percentage of embryonic viability of *seip-1(A185P)* is not affected in the day 2-3 (post L4 36-60 hours) animals. (C) DAPI staining of zygotic chromatin (yellow arrows) in *seip-1(A185P)* mutant embryos is indicative of a defective permeability barrier. (D) *seip-1(A185P)* exhibits enlarged LDs in the −1 to −3 oocytes marked by BODIPY. The insert panel shows 6x enlarged view of a single enlarged LD. (E) Quantification of enlarged LDs (diameter > 1.5 µm) in −1 to −3 oocytes of *seip-1(A185P)* animals. (F) The percentage of viable embryos was significantly rescued in the day 1 (post L4 0-36 hours) dihomo-γ-linolenic acid (FA20:3_n-6_) (PUFA)-fed *seip-1(A185P)* animals compared with control. In contrast, the percentage of viable embryos was reduced in the day 2-3 animals (36-60 post L4) fed with PUFAs. Scale bars are indicated in each panel. P-values: * < 0.05 (A, B, E); **** <0.0001 (B, G) (t-test).

## Discussion

Seipin is an ER-integrated protein that plays a key role in LD biology. Dysfunction of SEIPIN1 affects both adipocyte development and lipid storage in non-adipose tissues, which causes the various human diseases, such as lipodystrophy, neurological seipinopathies, and reproductive infertility. However, the detailed molecular and cellular mechanisms of these diseases are not well understood. How SEIPIN1 dysfunction affects lipid biosynthesis, metabolism, and utilization in building the lipid-rich organelles remain largely unknown. Here our studies report a comprehensive *in vivo* analysis of *C. elegans SEIPIN1*, *seip-1,* and suggest that the functional role of SEIP-1 is to maintain the lipid balance in *C. elegans* embryos, which is crucial for the proper formation of the permeability barrier of the eggshell that forms early in embryogenesis. Consistent with the observations in other SEIPIN-deficiency systems (human, mice, and yeast), loss of SEIP-1 was sufficient to impair lipid biosynthetic gene expression, lipid balance, and LD morphology. Furthermore, this study supports the notion that it is plausible to model BSCL2 or other SEIPIN-associated diseases in *C. elegans*. Importantly, the embryonic lethality and defective permeability barrier in *seip-1* mutants could be significantly rescued by the dietary supplementation of PUFAs (C18:3_n-6_and C20:3_n-6_), suggesting *C. elegans* is a powerful model system to screen potential genetic and pharmaceutical suppressors for SEIPIN1 diseases.

### *C. elegans SEIP-1* shares evolutionally conserved SEIPIN functions

A sole *SEIPIN* ortholog, *seip-1*, was identified in *C. elegans.* Although SEIP-1 shares 20% amino acid identity with both human SEIPIN1 and yeast Sei1p, a functional complementation assay indicated that *C. elegans seip-1* significantly suppresses the abnormal LD defects in the yeast *sei1* deficiency mutant. This finding is supported by a recent report in which expression of human *SEIPIN1* suppressed the abnormal LD size in a *C. elegans seip-1* mutant (Cao et al., 2019), suggesting *C. elegans seip-1* shares an evolutionarily-conserved function for LD homeostasis and morphology. The sub-cellular expression of endogenous SEIP-1::mScarlet indicated that SEIP-1 forms rings around a subset of LDs in different tissues, which is consistent with previous reports that SEIPIN is an ER-integrated protein and performs important roles during nascent LDs formation (Windpassinger et al., 2004). Additionally, *SEIPIN1* has important roles in controlling LD number and morphology (Fei et al., 2008; Szymanski et al., 2007). Our studies demonstrate that *seip-1* mutants cause the appearance of abnormally sized LDs in both adipose tissue (intestine) and non-adipose tissues such as oocytes and embryos. Interestingly, PUFA supplementation did not rescue the abnormal LD size defect, possibly suggesting that the primary function of Seipin, at least in *C. elegans*, may be in regulating lipid metabolism, and not LD biogenesis. The precise mechanisms of how SEIPIN regulates LD biogenesis and scaffolds the nascent LDs to the ER still remain obscure.

### Loss of SEIP-1 disrupted the regulation of fatty acid biosynthetic enzyme levels

Previous studies have identified anomalies in lipid homeostasis associated with SEIPIN mutations in humans. Consistent with previous studies in mice and humans showing that dysfunction of SEIPIN caused defects in fatty acid biosynthesis and metabolism (Chen et al., 2009; Jiang et al., 2014; Payne et al., 2008), our results suggest that *C. elegans seip-1* shares similar functions in maintaining fatty acid homeostasis. Fatty acids are building blocks of membrane and storage lipids. Fatty acids were known to affect cell type specific gene expression that impacts metabolic pathways, including lipid metabolism (Georgiadi and Kersten, 2012; Nakamura et al., 2004; Pegorier et al., 2004). Alternatively, fatty acids might be indirectly involved in regulating gene expression through their effects on specific enzyme-mediated pathways, membrane compositions, or signaling molecules. Although the precise role of SEIP-1 in regulating fatty acid metabolism has not been clear, it is key to understanding the *de novo* fatty acid synthesis pathway during embryogenesis. Equally important, as SEIP-1 is a crucial factor for fine-tuning LD size, is to determine whether or not LDs may contribute to cell or non-cell autonomous regulation of fatty acid metabolism.

### SEIP-1 is essential for permeability barrier formation in the eggshell

The *C. elegans* permeability barrier is a distinct envelope encasing the embryo, which is built at anaphase of meiosis II. This lipid-rich layer prevents the entry of potential toxic molecules and protects the embryos from osmotic damage. Previous studies identified fatty acid biosynthetic and modification enzymes, such as acetyl-CoA carboxylase POD-2 and fatty acid synthase FASN-1, as crucial for building the permeability barrier. Depletion of POD-2 and other synthetic enzymes in the germline impaired permeability barrier formation, suggesting that germline-derived fatty acids likely comprise the permeability barrier (Rappleye et al., 2003; Tagawa et al., 2001). Involvement of CYP-31A2/A3, DGTR-1, and PERM-1 suggests that these fatty acids might be utilized in the synthesis of lipid derivatives commonly stored within lipid droplets, such as triacylglycerol and sterol esters.

Transcriptional analysis from the mutant embryos suggest that SEIP-1 directly or indirectly influences the transcriptional expression of many lipid biosynthetic genes, including *perm-1* and *dgtr-1.* The reduced gene expression observed in the *seip-1* mutant embryos likely resulted in the deficit of lipid content, which is crucial to permeability barrier formation. Reduced PUFAs were also observed in these mutants; dietary supplementation with DGLA (C20:3_n-6_), significantly rescued the embryonic lethality and restored the defective permeability barrier. This finding suggests that DGLA may either be modified to the relevant lipid derivative found in the permeability barrier, or delivered alone to the extracellular matrix for building the permeability barrier. Combined with the high expression of *seip-1* at the sheath cells outside the ovulating oocyte, it is plausible to predict that SEIP-1 may deliver the LD with C20 long chain fatty acid derivative to the ovulating oocyte to prepare for permeability barrier formation.

### The *C. elegans* permeability barrier is a sensitive *in vivo* system in which to model human SEIPIN-associated disorders

SEIPIN is an important ER-associated protein that apparently has no enzymatic functions. Clinical reports indicate that either loss-of-function or gain-of-function mutations in the human *SEIPIN1* and *SEIPIN2* genes cause a variety of disorders. However, the molecular mechanism underlying all SEIPIN-associated disorders are still far from being understood. Disorders involving SEIPIN1 have been identified in different tissues, including both adipose tissue and non-adipose tissues. Recessive homozygous *SEIPIN1* loss-of-function mutations in humans cause BSCL type 2, a recessive disease. Affected individuals display a severe reduction of adipose tissue. Additionally, in these patients, *SEIPIN1* mutations also caused spermatozoid morphology defects and teratozoospermia. These phenotypes have not been well-studied in individuals with SEIPIN mutations however.

In this study, we found that *C. elegans seip-1* mutants displayed defects in LD size and also disrupted the formation of a lipid-rich layer of the protective eggshell. The *C. elegans seip-1* gene strongly expresses in sperm, and mutants cause severe reproductive defects, including smaller brood sizes and defects in eggshell formation. These phenotypes allowed us to investigate the role of *seip-1* in eggshell formation, a critical process necessary for embryogenesis and viability. The high expression in sperm is intriguing and we plan to investigate its function in a separate study. Though SEIPIN has been mostly associated with LD formation and morphology, it is curious that fatty acid biosynthetic gene expression in animals lacking SEIPIN are so dramatically altered. It suggests some feedback on lipid synthesis when LD formation is disrupted. In the studies presented here, the defects in permeability barrier formation, the readout of which is embryonic lethality, could be reversed with dietary supplementation of PUFAs. This observation suggests that a chemical suppressor screen could be beneficial.

Our observation that a BSCL2 patient-specific allele in the *C. elegans seip-1* gene displayed defects similar to our deletion alleles suggests that this allele is loss-of-function. These findings also demonstrate that *C. elegans* could be a good system in which to model lipodystrophies and seipinopathies. Decreased brood size and embryonic lethality are good readouts. Furthermore, a number of useful reagents exist to monitor the permeability barrier of the eggshell.

In summary, we demonstrate that *C. elegans* SEIPIN is required for the proper size of LDs, the proper expression of a number of lipid biosynthetic genes, and for the proper formation of the inner lipid-rich layer of the *C. elegans* eggshell. In this study, we chose to focus on the most striking phenotypes, the formation of the permeability barrier and fatty acid homeostasis, in which defects in each result in small brood sizes and embryonic lethality. The discovery of dietary rescue provides an avenue to study the molecular mechanism of SEIPIN *in vivo*. Further study will focus on understanding the molecular mechanisms of SEIPIN in regulating lipid biosynthesis during the reproductive process. We will attempt genetic suppressor screens to identify extragenic mutations that restore viability to *seip-1* mutant embryos. Likewise, we will attempt chemical suppressor screens that may help identify insightful approaches for future clinical therapy.

## Materials and Methods

### *C. elegans* strains used in this study

*C. elegans* strains were maintained with standard protocols. Strain information is listed in Table 1. AG547 and AG548 were created by crossing AG444 (*seip-1::mScarlet*) males with hermaphrodites containing *ocfIs2*[*pie-1p:GFP::SP12::pie-1 3’UTR + unc-119(+)*] and *pwIs23* [*vit-2::GFP*], respectively. We screened the F3 adults for the presence of the SEIP-1::mScarlet with GFP::SP12 and VIT-2::GFP transgenes, respectively, by microscopy. AG549 and AG550 were created by crossing OD344 males with hermaphrodites containing *seip-1(av109)* and *seip-1(tm4221)*, respectively. F3 adults expressing mCherry::CPG-2 and GFP::PH transgenes were genotyped by PCR for the *seip-1(av109)* and *seip-1(tm4221)* alleles. The *Saccharyomyces cerevisae* strain CWY3115 (*sei1Δ::HIS ERG6-mCherry::KAN BY4742*) was generated by a PCR-based integration method (Longtine et al., 1998). The yeast expression plasmids, *P_GPD_-GFP or pRS416-P_GPD_-GFP::seip-1* were transformed into the yeast cells. The transformed cells were grown in SCD media (0.67%YNB, amino acids, 2% glucose) to log phase (OD600=1.0) at 30°C. Another 24 hours extended growth allowed the cells to reach the stationary phase for visualization by microscopy.

### RNAi treatment

The RNAi feeding constructs were obtained from the Ahringer and Vidal libraries (Fraser et al., 2000; Rual et al., 2004). RNAi bacteria were grown until log phase was reached and spread on MYOB plates containing 1mM IPTG and 25 µg/ml carbenicillin and incubated overnight at 22°C. To silence the target genes, mid-L4 hermaphrodites were picked onto plates with the IPTG-induced bacteria. Animals were grown on RNAi plates at 20°C.

### CRISPR design

We used the Bristol N2 strain as the wild type for CRISPR/Cas9 editing. The gene-specific 20 nucleotide sequences for crRNA synthesis were selected with help of a guide RNA design checker from Integrated DNA Technologies (IDT) (https://www.idtdna.com) and were ordered as 20 nmol or 4 nmol products from Dharmacon (https://dharmacon.horizondiscovery.com), along with tracRNA. Repair template design followed the standard protocols (Paix et al., 2015; Vicencio et al., 2019). Approximately 30 young gravid animals were injected with the prepared CRISPR/Cas9 injection mix as described in the literature (Paix et al., 2015). *seip-1(av109)* was generated by CRISPR/Cas9 mixes that contained two guide RNAs at flanking regions of *seip-1* coding regions. Heterozygous *seip-1* deletion animals were first screened by PCR and then homozygosed in subsequent generations. mScarlet insertions at the *seip-1* C-terminus were performed by Nested CRISPR (Vicencio et al., 2019). Homozygous *nest-1* edited animals were confirmed by PCR and restriction enzyme digestion and selected for the secondary CRISPR/Cas9 editing. Full-length mScarlet insertion animals were screened by PCR and visualized by fluorescence microscopy. All homozygous animals edited by CRISPR/Cas9 were confirmed by Sanger sequencing (Eurofins). The detailed sequence information of the repair template and guide RNAs are listed in Table 2. Two *SEIP-1* (*R01B10.6*) deletion strains were custom made by Knudra Transgenics (currently Nemamatrix, Inc) (Murray, UT) using three proprietary vectors, the locus-targeting guide RNA pNU1244, the locus-targeting guide RNA PNU1245, and the donor homology plasmid pNU1246. Along with standard fluorescence markers and Cas9 plasmid (pNU792), the DNA mix is injected into gonads of the N2 strain, and the injected animals were screened for resistance to 0.25 mg/ml hygromycin followed by a heat shock counter screen to remove animals bearing extrachromosomal arrays. Two verified transgenic lines received from Knudra were backcrossed with N2 four times to obtain the *seip-1(cwc1)* and *seip-1(cwc2)* strains that were confirmed by PCR.

### Brood size determinations and embryonic viability assays

Single mid-L4 hermaphrodites were picked onto 35 mm MYOB plates seeded with 10 µl of OP50 bacteria and allowed to lay eggs for 36 hours (plate one contains the brood from 0-36 hours post mid-L4). The same hermaphrodite was moved to a new 35 mm MYOB plate to lay eggs for another 24 hours and was removed from the plate (this plate contains the brood from 36-60 hours post mid-L4). Twenty-four hours after removing the moms, only fertilized embryos and larvae were counted to determine the brood size. Brood sizes were determined at 36 hours and 60 hours. Percentage of embryonic viability= (the number of hatched larva / the total number of hatched and unhatched animals) *100%. The embryonic lethality of *seip-1*(*cwc1* and *cwc2*) deletion strains was much higher (∼75-80%) than that of the *av109* deletion allele. It is possible that the difference in lethality observed is due in part by the culturing conditions used between two laboratories. The *cwc* strains were grown on standard NGM plates while the *av109* allele was grown on MYOB plates. They differ in pH and some salts. In addition, we occasionally observed adaptive rescue of lethality (or the acquisition of spontaneous suppressors) after maintaining homozygous strains over a period of a few months. Thus, the *cwc1* and *cwc2* strains were revived from frozen stocks every three months. Such spontaneous suppressors have been observed for unrelated temperature-sensitive embryonic lethal mutations (K. O’Connell, personal communication).

### BODIPY 493/503 staining

BODIPY 493/503 (Invitrogen # D3922) was dissolved in 100% DMSO to 1 mg/ml. BODIPY stock was diluted by M9 to 6.7 µg/ml BODIPY (final concentration of DMSO was 0.8%) as the working stock. Hermaphrodites were washed in M9 three times and incubated in 6.7 µg/ml BODIPY for 20 minutes and washed again in M9 at least three times. All washes and incubations were performed in a concavity slide (ThermoFisher, # S99369). The stained hermaphrodites were anesthetized with 0.1% tricaine and 0.01% tetramisole in M9 buffer for 15-30 minutes. The anesthetized animals were then transferred to 5% agarose pads for imaging. Image acquisition was captured by a Nikon 60 X 1.2 NA water objective with 0.5 µm z-step size.

### DAPI staining of embryos

Hermaphrodites were washed in M9 three times and dissected with 23 G x 3/4” needles. Different stage embryos were transferred to a hanging drop chamber filled with Blastomere Culture Medium (BCM) (Edgar and Goldstein, 2012). The Blastomere Culture Medium was made fresh with 5 µg/ml DAPI. The hanging drop chamber was sealed by molten Vaseline before imaging. Image acquisition was captured by a Nikon 60 X 1.2 NA water objective with 1 µm z-step size.

### FM4-64 staining of embryos

To check eggshell integrity, embryos were dissected from both wildtype and *seip-1(cwc1)* animals in 5 µl egg buffer (4 mM HEPES pH7.4, 94 mM NaCl, 32 mM KCl, 2.7 mM CaCl_2_ and 2.7 mM MgCl_2_) containing 6.6 µM FM4-64 (ThermoFisher, # T3166). Samples were mounted in a chamber circled by vacuum grease on a 2% agar pad (15 x 15 mm) for imaging. The images were acquired on a ZEISS Z1 fluorescence microscopy system that uses an EC Plan-Neofluar 40X objective lens (NA0.75), an EC Plan-Neofluar M27 20X objective lens (NA0.5), a mCherry filter set (Ex 540-552nm, Em 575-640nm) and a Zeiss AxioChem 506 CCD camera. The images were acquired and analyzed by ZEISS ZEN software.

### Live imaging of ovulation

For imaging SEIP-1::mScarlet expression during ovulation, animals were immobilized on 4% agar pads with anesthetic (0.1% tricaine and 0.01% tetramisole in M9 buffer). DIC and mScarlet image acquisition were captured by a Nikon 60 X 1.2 NA water objective with 1-2 µm z-step size; 10-15 z planes were captured. Time interval for ovulation imaging is every 30 seconds, and duration of imaging is 10-15 minutes.

### Imaging of yolk proteins

Day 1 adults (24 hours post mid-L4) of AG548 were immobilized on 4% agar pads with anesthetic (0.1% tricaine and 0.01% tetramisole in M9 buffer). The acquisition of both DIC and 561 nm images were performed by our confocal imaging system (see below) with a Nikon 60 X 1.2 NA water objective. 10-20 z planes were captured with 1 µm z-step size. Images were generated by custom Fiji code using Image>Stacks>Z Project.

### Imaging of LDs in yeast cells

To image yeast cells, yeast cells were cultured in SCD medium to stationary phase and were imaged by a ZEISS Z1 fluorescence microscopy system that uses a ZEISS Plan-Apochromat 100X objective lens (NA 1.4), a GFP filter set (Ex 450-490 nm, Em 500-550 nm), a mCherry filter set (Ex 540-552 nm, Em 575-640 nm), and a Zeiss AxioChem 506 CCD camera. The images were acquired and analyzed by ZEISS ZEN software.

### PUFA dietary supplementation

PUFA dietary supplementation was modified from a previous report (Yang et al., 2006). Linoleic acid (C18:2_n-6_, Sigma # L1376), γ-linolenic acid (C18:3_n-6_, Sigma # L2378), Dihomo-γ-linolenic acid (C20:3_n-6_, Cayman Chemical # 90230) were added to NGM or MYOB medium to a final concentration of 300 µM. Single mid-L4 hermaphrodites were picked onto 35 mm PUFA-supplemented MYOB and control MYOB plates seeded with 10 µl of OP50 bacteria and allowed to lay eggs for 36 hours (plate one contains the brood from 0-36 hours post mid-L4). The same hermaphrodite was moved to a new 35 mm MYOB plate to lay eggs for another 24 hours and were removed from the plate (this plate contains the brood from 36-60 hours post mid-L4). Twenty-four hours after removing the moms, only fertilized embryos and larvae were counted to determine the brood size. Brood sizes were determined at 36 hours and 60 hours. Percentage of embryonic viability= (the number of hatched larva / the total number of hatched and unhatched animals) *100%. For embryo imaging, single mid-L4 hermaphrodites were allowed to grow on PUFA-supplemented MYOB for 24 hours. Then these gravid adults were dissected in 5 µl of 0.7X egg buffer or BCM containing either 6.6 µM FM4-64 or 5 µg/ml DAPI. The embryos were mounted in a hanging drop chamber and images were acquired as described above.

### Electron microscopy

∼60 gravid hermaphrodites (60 h after L1) were picked (10∼12 animals per grid) and mixed with fresh yeast, which were immediately subjected to cryofixation by high-pressure freezing with a high-pressure freezer EM HPM100 (Leica, Wetzlar, Germany). AFS2 EM (Leica, Wetzler, Germany) was used for the subsequent freeze substitution and low temperature embedding. Freeze substitution was performed in 2% osmium tetroxide and 0.1% uranyl acetate in acetone following the program of −90°C for 16 hrs, −85°C for 50 hrs, −50°C for 10 hrs, −20°C for 10 hrs, and 0°C for 6 hrs. Temperature increased by a rate of 2.5°C / hr for each time slope. The samples were then washed in acetone and subjected to infiltration with 5%, 10%, 25%, 50%, 75%, and final 100% Spurr resin. After polymerization at 60°C for 48 hrs, the blocks were cut into 70-90 nm ultrathin sections on the ultramicrotome (EM UC6; Leica) with a diamond knife (Ultra 45°, DiATOME; Biel, Switzerland). Sections were placed on 0.4% Formvarcoated slot grids (EMS) and imaged at 120 kV on a transmission electron microscope (Tecnai G2 Spirit TWIN; FEI, Hillsboro, OR) with a digital CCD camera (832 Orius SC1000B; Gatan, Pleasanton, CA). The images were processed and analyzed by Photoshop software (CS4, Adobe).

### Microarray

Embryos were isolated from gravid adults by hypochlorite disruption (0.5N NaOH and 1% NaOCl) followed by M9 washes. After overnight incubations in M9 without food, synchronized L1 were then seeded on a 5.5cm NGM plate (∼150 L1 /plate). The synchronized young adult (52 hrs post L1 feeding) were harvested and washed by M9 buffer and pelleted. Embryos were isolated by hypochlorite disruption from young gravid adult (52 hrs after L1), washed with M9, and pelleted. To extract total RNA, synchronized young adult or embryos derived from ∼40,000 young adults were resuspended in 500 µl TRIzol reagent followed by flash freezing in liquid nitrogen. Samples were thawed at 42°C and frozen in liquid nitrogen and the cycle was repeated 6-7 times. 100 µl chloroform was mixed with sample and spun at 12,000g for 15 minutes at 4°C. The corresponding aqueous layer was harvested and mixed with equal volume of 70% ethanol. RNeasy Mini kits (Qiagen) were used to obtain pure RNA for microarray analysis. A 0.2 µg of total RNA was subjected to Cy3 labeling by *in vitro* transcription with use of Low Input Quick-Amp Labeling kit (Agilent Technologies, USA). Subsequently, a 1.65 µg of Cy3-labled cRNA was incubated at 60°C for 30 minutes, followed by hybridization to the Agilent *C. elegans* V2 4*44K Microarray chip (Agilent Technologies, USA) at 65°C for 17 hrs. After washing and drying, the microarray chip was scanned with the microarray scanner (Agilent Technologies, USA) at 535 nm for Cy3. The array image was read by Feature Extraction software version 10.7.1.1, and data was subjected to further analysis by GeneSpring software (Agilent). Average value of N2 sample was used as the control; a statistics filter was set to select for *seip-1*/N2 gene expression at a fold change >=1.5 and p-value <=0.05. Genes that passed these criteria were subjected to KEGG pathway annotation using DAVID Bioinformatics Resources 6.8 (https://david.ncifcrf.gov/home.jsp).

### Quantitative qPCR

*C. elegans* animals were first synchronized by the hypochlorite disruption method as described above. Expression levels of SEIP-1 were carried out in whole animal or tissues dissected from animals. RNA prepared immediately (0 hour) or 5 hour after embryos were dissected from animals were compared. Embryos isolated from ∼40,000 young adult (52 hrs after L1) were resuspend in 500 µl TRIzol Reagent followed by flash freezing in nitrogen liquid. Samples were thawed at 42°C and frozen in liquid nitrogen and the cycle was repeated 6-7 times. 100 µl chloroform was mixed with sample and spun at 12,000g for 15 minutes at 4°C. The corresponding aqueous layer was harvested and mixed with equal volume of 70% ethanol. RNeasy Mini kits (Qiagen) were used to extract pure RNA. ∼15 µg RNA in nuclease-free water was treated by DNase I. 2 µg DNaseI-treated RNA was used as the template to synthesize cDNAs with the SuperScript III First Strand system (Thermo Fisher Scientific). The real-time PCR reactions were performed by the Applied Biosystems™ QuantStudio™ 12K Flex system with power SYBR Green PCR master mix (Thermo Fisher Scientific). The thermal cycling conditions followed an initial denaturation step at 95°C for 10 min, 40 cycles at 95°C for 10 sec, and 60°C for 1 min followed by a continuous melt curve stage at 95°C for 15 sec, 60°C for 1 min, 95°C for 15 sec. Three independent experiments were replicated for each target gene. The ΔCT value was determined by QuantStudio™ 12K Flex software (Thermo Fisher Scientific). Gene expression levels were calculated as 2^[−ΔΔC(T)], which was normalized to the expression of the *act-1* gene. Statistical analyses were calculated based on four biological replicates and shown as average +/− standard deviation.

## Acknowledgments

We thank the *Caenorhabditis* Genetics Center, which is funded by National Institutes of Health Office of Research Infrastructure Programs (P40OD010440), for providing strains for this study. We thank Dr. Shohei Mitani for sharing the FX14734 [*seip-1(tm4221)/hT1 V*] strain. We thank Dr. Orna Cohen-Fix for generously sharing the GFP::SP-12 strain. We are grateful to the members of the Golden laboratory: Dr. Peter Kropp, Dr. Tao Cai, Rosie Bauer, Isabella Zafra for productive discussions and preparing reagents. We thank Dr. Rey-Huei Chen (IMB, Academia Sinica) for sharing lab resources and improving the manuscript. We are grateful to Tzu-Han Hsu (IMB, Academia Sinica) for advice on EM, Ji-Ying Huang and Mei-Jane Fang at the Cell Biology Core Lab (IPMB, Academia Sinica) for advice on microscopy and qPCR analysis, Shu-Jen Chou at the Genomic Technology Core Lab (IPMB, Academia Sinica) for help with microarray, and the IPMB Bioinformatic Core Lab for consulting on genomic analysis. We especially thank Dr. William Prinz and Dr. Alexandre Toulmay for critical inputs on the project and feedback on the manuscript. We thank all members of the NIH and Baltimore Worm Club for providing feedback and suggestions to our investigations.

---

**Supplemental Figure 1. Expression of *C. elegans seip-1* rescues abnormal LD morphology defects in yeast *sei1*Δ mutants.**

(A-D) Plasmid with control GFP (green, A, C) was transformed and expressed in the yeast *sei1Δ* strain expressing the LD marker Erg6p-mCherry (red, B, C). Red arrows indicate supersized LDs in the *sei1Δ* cells. (E-H) Plasmid with *GFP-SEIP-1* gene (green, E, G) was transformed and expressed in the yeast *sei1Δ* strain expressing the LD marker Erg6p-mCherry. GFP-SEIP-1p was observed adjacent to or surrounding the LDs labeled by Erg6p-mCherry (blue arrows, E-G). (I-L) Co-localization of GFP-SEIP-1p with ER marker Sec63p-mCherry on the LDs (pink arrows) was observed in the *sei1Δ* cells. (M) Quantification of LD morphology in the *sei1Δ* cells expressing either control GFP or GFP-SEIP-1p. n= the number of quantified cells in the study. Scale bars are indicated in each panel.

**Supplemental Figure 2. *seip-1* expression pattern in *C. elegans***

(A) Expression levels of SEIP-1 in whole animal or tissues dissected from animals. qRT-PCR was performed using RNA prepared immediately (0 hour) or 5 hours after embryos were dissected from animals were compared. (B-G) SEIP-1::mScarlet (magenta) expresses in a variety of cell types, including embryos (B, B’), pharynx (C, C’), tail and rectum (D, D’), hypodermal cells (E, E’), intestinal cells (F-G). The small panels in G represent enlarged views of the yellow square. SEIP-1::mScarlet labeled LD is indicated by yellow arrows. Small top panel represents the merge image, middle panel is SEIP-1::mScarlet only, bottom panel is BODIPY staining image only. (H-J) SEIP-1::mScarlet also expresses on the sperm (yellow box represents the constricted spermatheca and white arrows mark the individual sperm). Scale bars are indicated in each panel.

**Supplemental Figure 3. Verification of CRISPR/Cas9 generated deletions in *seip-1* knockout animals.**

(A-C) Schematic of all *seip-1* knockout alleles and the position of the genotyping primers (F1, R1 and R2) used in this study. (B) *seip-1(cwc1)* and *seip-1(cwc2)* lines are shown as a replacement of the *seip-1* gene with an expression cassette of a hygromycin B resistance gene (*hygR*). (C) *seip-1(av109)* was generated by CRISPR to delete the whole *seip-1* gene. The positions of the genotyping primers (F2, R3 and R4) for verifying the knockout alleles are marked by the pink lines that flank the *seip-1* coding exons. Arrows represent the primer direction from 5’ to 3’ (A-C). (D-F) Representative PCR gel from genotyping single animals for *seip-1(cwc1)* or *seip-1(cwc2)* candidates. Two pairs of primers [one pair (F1 and R1) for wt (A); the other pair (F1 and R2) for KO insertion (B)] were used to genotype *seip-1(cwc1)* and *seip-1(cwc2)* candidates. In wild type, PCR product was able to be amplified by wt primers (D), but not by KO primers. Homozygous knockout was only able to be amplified by KO primers but not by wt primers (E, F). (G) Three primers (two flanking primers F2 and R4 locate outside the deleted region and an internal primer R3) were used to genotype for homozygosity of candidate *seip-1(av109)* deletion animals. Amplicon size of a homozygous deletion with both flanking primers is 422 bp. In wild type, the PCR product amplified by one flanking primer and the internal primer is 877 bp. Heterozygous animals contain both PCR products. (H) Expression of the *seip-1* rescuing fosmids and plasmids in *seip-1(cwc1)* mutant significantly rescues the embryonic lethality. **** <0.0001 (H) (t-test).

**Supplemental Figure 4. Deletion of the *seip-1* gene disrupted permeability barrier formation.**

(A, B) DIC images show that multicellular *seip-1(cwc1)* mutant embryos (B) displayed a shrunken and misshapen morphology compared with wildtype embryos (A). (C) Diagram of deleted region in the *seip-1(tm4221)* mutant. (D, E) The penetration of both DAPI (D) and mCherry::CPG-2 (E) indicated that *seip-1(tm4221)* embryos also were defective for permeability barrier formation. Zygotic chromatin was stained with DAPI in *seip-1(tm4221)* (yellow arrowheads D). mCherry::CPG-2 (magenta) penetrates inside the entire space between eggshell and the embryo surface (E) (white arrowheads) (n=number of embryos with mCherry::CPG-2 penetration / number of embryos imaged). Scale bars are indicated in each panel.

**Supplemental Figure 5. Depletion of the *seip-1* gene alters lipid droplets size in the oocytes and intestine.**

(A) Quantification of total enlarged LDs (diameter > 1.5 µm) in the −1 to −3 oocytes of both wildtype and *seip-1(cwc1)* animals. (B) Transmission electron micrographs of high-pressure frozen animals displayed LDs in both wildtype and *seip-1(cwc1)* intestinal cells. Red asterisks mark the normal size LDs in wild type, red arrows indicate smaller size LDs, blue asterisks indicate enlarged LDs. Inserts are 6X amplified images of the inlaid yellow square regions. Scale bar is indicated.

**Supplemental Figure 6. Global gene expression of *seip-1* mutants were analyzed by the microarray**

(A) Heatmap shows the variation of 20,221 genes expressed between *seip-1(cwc1)* mutants and wild type. The data were collected from two independent experiments. Mis-regulated gene expression were found in *seip-1(cwc1)* embryos compared to wild type. (B) The genes with differential expressions between *seip-1(cwc1)* mutant and wild-type embryos were identified by KEGG (Kyoto Encyclopedia of Genes and Genomes) pathway enrichment analyses. Metabolic pathways genes were significantly affected when the *seip-1* gene was deleted.

**Video S1. SEIP-1 expression pattern during ovulation**

Ovulation imaged in the genome-edited animals expressing SEIP-1::mScarlet (magenta). arrows in left and right panels indicate SEIP-1::mScarlet expression in the fifth gonadal sheath cell until the oocyte completed the ovulation. Left panel indicates the mScarlet (magenta) channel only, middle channel indicates DIC only, right channel indicates the merged image of mScarlet and DIC. Images are z-max with 10 z planes taken every 30 seconds with 1 µm step size. Timing is indicated in lower right of the left panel. Playback rate is 2 frames/second. Scale bar is indicated in the left panel.

## References

Benenati, G., Penkov, S., Muller-Reichert, T., Entchev, E.V., and Kurzchalia, T.V. (2009). Two cytochrome P450s in Caenorhabditis elegans are essential for the organization of eggshell, correct execution of meiosis and the polarization of embryo. Mechanisms of Development 126, 382–393.

Cao, Z., Hao, Y., Fung, C.W., Lee, Y.Y., Wang, P.F., Li, X.S., Xie, K., Lam, W.J., Qiu, Y.F., Tang, B.Z., et al. (2019). Dietary fatty acids promote lipid droplet diversity through seipin enrichment in an ER subdomain. Nature Communications 10.

Cartwright, B.R., and Goodman, J.M. (2012). Thematic Review Series: Lipid Droplet Synthesis and Metabolism: from Yeast to Man Seipin: from human disease to molecular mechanism. J Lipid Res 53, 1042–1055.

Chen, W., Yechoor, V.K., Chang, B.H., Li, M.V., March, K.L., and Chan, L. (2009). The human lipodystrophy gene product Berardinelli-Seip congenital lipodystrophy 2/seipin plays a key role in adipocyte differentiation. Endocrinology 150, 4552–4561.

Edgar, L.G., and Goldstein, B. (2012). Culture and Manipulation of Embryonic Cells. Method Cell Biol 107, 153–175.

El Zowalaty, A.E., Baumann, C., Li, R., Chen, W., De La Fuente, R., and Ye, X. (2015). Seipin deficiency increases chromocenter fragmentation and disrupts acrosome formation leading to male infertility. Cell Death Dis 6, e1817.

Fei, W.H., Li, H., Shui, G.H., Kapterian, T.S., Bielby, C., Du, X.M., Brown, A.J., Li, P., Wenk, M.R., Liu, P.S., et al. (2011). Molecular characterization of seipin and its mutants: implications for seipin in triacylglycerol synthesis. J Lipid Res 52, 2136–2147.

Fei, W.H., Shui, G.H., Gaeta, B., Du, X.M., Kuerschner, L., Li, P., Brown, A.J., Wenk, M.R., Parton, R.G., and Yang, H.Y. (2008). Fld1p, a functional homologue of human seipin, regulates the size of lipid droplets in yeast. J Cell Biol 180, 473–482.

Fraser, A.G., Kamath, R.S., Zipperlen, P., Martinez-Campos, M., Sohrmann, M., and Ahringer, J. (2000). Functional genomic analysis of C. elegans chromosome I by systematic RNA interference. Nature 408, 325–330.

Garg, A. (2004). Acquired and inherited lipodystrophies. N Engl J Med 350, 1220–1234.

Georgiadi, A., and Kersten, S. (2012). Mechanisms of gene regulation by fatty acids. Adv Nutr 3, 127–134.

Golden, A., Liu, J., and Cohen-Fix, O. (2009). Inactivation of the C. elegans lipin homolog leads to ER disorganization and to defects in the breakdown and reassembly of the nuclear envelope. J Cell Sci 122, 1970–1978.

Gonzalez, D.P., Lamb, H.V., Partida, D., Wilson, Z.T., Harrison, M.C., Prieto, J.A., Moresco, J.J., Diedrich, J.K., Yates, J.R., 3rd, and Olson, S.K. (2018). CBD-1 organizes two independent complexes required for eggshell vitelline layer formation and egg activation in C. elegans. Dev Biol 442, 288–300.

Grippa, A., Buxo, L., Mora, G., Funaya, C., Idrissi, F.Z., Mancuso, F., Gomez, R., Muntanya, J., Sabido, E., and Carvalho, P. (2015). The seipin complex Fld1/Ldb16 stabilizes ER-lipid droplet contact sites. J Cell Biol 211, 829–844.

Gunsalus, K.C., Ge, H., Schetter, A.J., Goldberg, D.S., Han, J.D.J., Hao, T., Berriz, G.F., Bertin, N., Huang, J., Chuang, L.S., et al. (2005). Predictive models of molecular machines involved in Caenorhabditis elegans early embryogenesis. Nature 436, 861–865.

Hall, D.H., Winfrey, V.P., Blaeuer, G., Hoffman, L.H., Furuta, T., Rose, K.L., Hobert, O., and Greenstein, D. (1999). Ultrastructural features of the adult hermaphrodite gonad of Caenorhabditis elegans: Relations between the germ line and soma. Developmental Biology 212, 101–123.

Ito, D., Fujisawa, T., Iida, H., and Suzuki, N. (2008). Characterization of seipin/BSCL2, a protein associated with spastic paraplegia 17. Neurobiol Dis 31, 266–277.

Jiang, M., Gao, M., Wu, C., He, H., Guo, X., Zhou, Z., Yang, H., Xiao, X., Liu, G., and Sha, J. (2014). Lack of testicular seipin causes teratozoospermia syndrome in men. Proc Natl Acad Sci U S A 111, 7054–7059.

Johnston, W.L., and Dennis, J.W. (2012). The eggshell in the C. elegans oocyte-to-embryo transition. Genesis 50, 333–349.

Kanehisa, M., Sato, Y., Furumichi, M., Morishima, K., and Tanabe, M. (2019). New approach for understanding genome variations in KEGG. Nucleic Acids Res 47, D590–D595.

Kimble, J., and Sharrock, W.J. (1983). Tissue-Specific Synthesis of Yolk Proteins in Caenorhabditis-Elegans. Developmental Biology 96, 189–196.

Li, S.W., Xu, S.B., Ma, Y.L., Wu, S., Feng, Y., Cui, Q.P., Chen, L.F., Zhou, S., Kong, Y.Y., Zhang, X.Y., et al. (2016). A Genetic Screen for Mutants with Supersized Lipid Droplets in Caenorhabditis elegans. G3-Genes Genomes Genetics 6, 2407–2419.

Longtine, M.S., McKenzie, A., 3rd, Demarini, D.J., Shah, N.G., Wach, A., Brachat, A., Philippsen, P., and Pringle, J.R. (1998). Additional modules for versatile and economical PCR-based gene deletion and modification in Saccharomyces cerevisiae. Yeast 14, 953–961.

Magre, J., Delepine, M., Khallouf, E., Gedde-Dahl, T., Van Maldergem, L., Sobel, E., Papp, J., Meier, M., Megarbane, A., Lathrop, M., et al. (2001). Identification of the gene altered in Berardinelli-Seip congenital lipodystrophy on chromosome 11q13. Nature Genetics 28, 365–370.

Mak, H.Y. (2012). Lipid droplets as fat storage organelles in Caenorhabditis elegans: Thematic Review Series: Lipid Droplet Synthesis and Metabolism: from Yeast to Man. J Lipid Res 53, 28–33.

Nakamura, M.T., Cheon, Y., Li, Y., and Nara, T.Y. (2004). Mechanisms of regulation of gene expression by fatty acids. Lipids 39, 1077–1083.

Olson, S.K., Greenan, G., Desai, A., Muller-Reichert, T., and Oegema, K. (2012). Hierarchical assembly of the eggshell and permeability barrier in C. elegans. J Cell Biol 198, 731–748.

Paix, A., Folkmann, A., Rasoloson, D., and Seydoux, G. (2015). High Efficiency, Homology-Directed Genome Editing in Caenorhabditis elegans Using CRISPR-Cas9 Ribonucleoprotein Complexes. Genetics 201, 47-+.

Paupard, M.C., Miller, A., Grant, B., Hirsh, D., and Hall, D.H. (2001). Immuno-EM localization of GFP-tagged yolk proteins in C-elegans using microwave fixation. Journal of Histochemistry & Cytochemistry 49, 949–956.

Payne, V.A., Grimsey, N., Tuthill, A., Virtue, S., Gray, S.L., Nora, E.D., Semple, R.K., O’Rahilly, S., and Rochford, J.J. (2008). The human lipodystrophy gene BSCL2/Seipin may be essential for normal adipocyte differentiation. Diabetes 57, 2055–2060.

Pegorier, J.P., Le May, C., and Girard, J. (2004). Control of gene expression by fatty acids. J Nutr 134, 2444S–2449S.

Poteryaev, D., Squirrell, J.M., Campbell, J.M., White, J.G., and Spang, A. (2005). Involvement of the actin cytoskeleton and homotypic membrane fusion in ER dynamics in Caenorhabditis elegans. Molecular Biology of the Cell 16, 2139–2153.

Radman, I., Greiss, S., and Chin, J.W. (2013). Efficient and rapid C. elegans transgenesis by bombardment and hygromycin B selection. PLoS One 8, e76019.

Rappleye, C.A., Tagawa, A., Le Bot, N., Ahringer, J., and Aroian, R.V. (2003). Involvement of fatty acid pathways and cortical interaction of the pronuclear complex in Caenorhabditis elegans embryonic polarity. BMC Dev Biol 3, 8.

Rual, J.F., Ceron, J., Koreth, J., Hao, T., Nicot, A.S., Hirozane-Kishikawa, T., Vandenhaute, J., Orkin, S.H., Hill, D.E., van den Heuvel, S., et al. (2004). Toward improving Caenorhabditis elegans phenome mapping with an ORFeome-based RNAi library. Genome Res 14, 2162–2168.

Salo, V.T., Belevich, I., Li, S.Q., Karhinen, L., Vihinen, H., Vigouroux, C., Magre, J., Thiele, C., Holtta-Vuori, M., Jokitalo, E., et al. (2016). Seipin regulates ER-lipid droplet contacts and cargo delivery. Embo Journal 35, 2699–2716.

Schneider, W.J. (1996). Vitellogenin receptors: Oocyte-specific members of the low-density lipoprotein receptor supergene family. International Review of Cytology - a Survey of Cell Biology, Vol 166 166, 103–137.

Shi, X., Li, J., Zou, X.J., Greggain, J., Rodkaer, S.V., Faergeman, N.J., Liang, B., and Watts, J.L. (2013). Regulation of lipid droplet size and phospholipid composition by stearoyl-CoA desaturase. J Lipid Res 54, 2504–2514.

Shi, X., Tarazona, P., Brock, T.J., Browse, J., Feussner, I., and Watts, J.L. (2016). A Caenorhabditis elegans model for ether lipid biosynthesis and function. Journal of lipid research 57, 265–275.

Simha, V., and Garg, A. (2003). Phenotypic heterogeneity in body fat distribution in patients with congenital generalized lipodystrophy caused by mutations in the AGPAT2 or seipin genes. J Clin Endocrinol Metab 88, 5433–5437.

Stein, K.K., and Golden, A. (2018). The C. elegans eggshell. WormBook 2018, 1–36.

Sturmey, R.G., Reis, A., Leese, H.J., and McEvoy, T.G. (2009). Role of fatty acids in energy provision during oocyte maturation and early embryo development. Reproduction in domestic animals = Zuchthygiene 44 Suppl 3, 50–58.

Szymanski, K.M., Binns, D., Bartz, R., Grishin, N.V., Li, W.P., Agarwal, A.K., Garg, A., Anderson, R.G.W., and Goodman, J.M. (2007). The lipodystrophy protein seipin is found at endoplasmic reticulum lipid droplet junctions and is important for droplet morphology. Proceedings of the National Academy of Sciences of the United States of America 104, 20890–20895.

Tagawa, A., Rappleye, C.A., and Aroian, R.V. (2001). Pod-2, along with pod-1, defines a new class of genes required for polarity in the early Caenorhabditis elegans embryo. Dev Biol 233, 412–424.

Vicencio, J., Martinez-Fernandez, C., Serrat, X., and Ceron, J. (2019). Efficient Generation of Endogenous Fluorescent Reporters by Nested CRISPR in Caenorhabditis elegans. Genetics 211, 1143–1154.

Vrablik, T.L., Petyuk, V.A., Larson, E.M., Smith, R.D., and Watts, J.L. (2015). Lipidomic and proteomic analysis of Caenorhabditis elegans lipid droplets and identification of ACS-4 as a lipid droplet-associated protein. Biochimica Et Biophysica Acta-Molecular and Cell Biology of Lipids 1851, 1337–1345.

Walther, T.C., Chung, J., and Farese, R.V., Jr. (2017). Lipid Droplet Biogenesis. Annu Rev Cell Dev Biol 33, 491–510.

Wang, H.J., Becuwe, M., Housden, B.E., Chitraju, C., Porras, A.J., Graham, M.M., Liu, X.R.N., Thiam, A.R., Savage, D.B., Agarwal, A.K., et al. (2016a). Seipin is required for converting nascent to mature lipid droplets. Elife 5.

Wang, H.Z., Jiang, X., Wu, J.Y., Zhang, L.Q., Huang, J.F., Zhang, Y.R., Zou, X.J., and Liang, B. (2016b). Iron Overload Coordinately Promotes Ferritin Expression and Fat Accumulation in Caenorhabditis elegans. Genetics 203, 241-+.

Watts, J.L., and Ristow, M. (2017). Lipid and Carbohydrate Metabolism in Caenorhabditis elegans. Genetics 207, 413–446.

Watts, J.S., Morton, D.G., Kemphues, K.J., and Watts, J.L. (2018). The biotin-ligating protein BPL-1 is critical for lipid biosynthesis and polarization of the Caenorhabditis elegans embryo. Journal of Biological Chemistry 293, 610–622.

Windpassinger, C., Auer-Grumbach, M., Irobi, J., Patel, H., Petek, E., Horl, G., Malli, R., Reed, J.A., Dierick, I., Verpoorten, N., et al. (2004). Heterozygous missense mutations in BSCL2 are associated with distal hereditary motor neuropathy and Silver syndrome. Nat Genet 36, 271–276.

Wolinski, H., Kolb, D., Hermann, S., Koning, R.I., and Kohlwein, S.D. (2011). A role for seipin in lipid droplet dynamics and inheritance in yeast. Journal of Cell Science 124, 3894–3904.

Xu, N.Y., Zhang, S.B.O., Cole, R.A., McKinney, S.A., Guo, F.L., Haas, J.T., Bobba, S., Farese, R.V., and Mak, H.Y. (2012). The FATP1-DGAT2 complex facilitates lipid droplet expansion at the ER-lipid droplet interface. J Cell Biol 198, 895–911.

Yang, F., Vought, B.W., Satterlee, J.S., Walker, A.K., Jim Sun, Z.Y., Watts, J.L., DeBeaumont, R., Saito, R.M., Hyberts, S.G., Yang, S., et al. (2006). An ARC/Mediator subunit required for SREBP control of cholesterol and lipid homeostasis. Nature 442, 700–704.

Zhang, P., Na, H.M., Liu, Z.L., Zhang, S.Y., Xue, P., Chen, Y., Pu, J., Peng, G., Huang, X., Yang, F.Q., et al. (2012). Proteomic Study and Marker Protein Identification of Caenorhabditis elegans Lipid Droplets. Molecular & Cellular Proteomics 11, 317–328.

Zowalaty, A.E.E., and Ye, X. (2017). Seipin deficiency leads to defective parturition in mice. Biol Reprod 97, 378–386.

