## Supplemental Figures for "Dietary supplementation with PUFAs rescues the eggshell defects caused by *seipin* mutations in *C. elegans*"

### Supplemental Figure 1

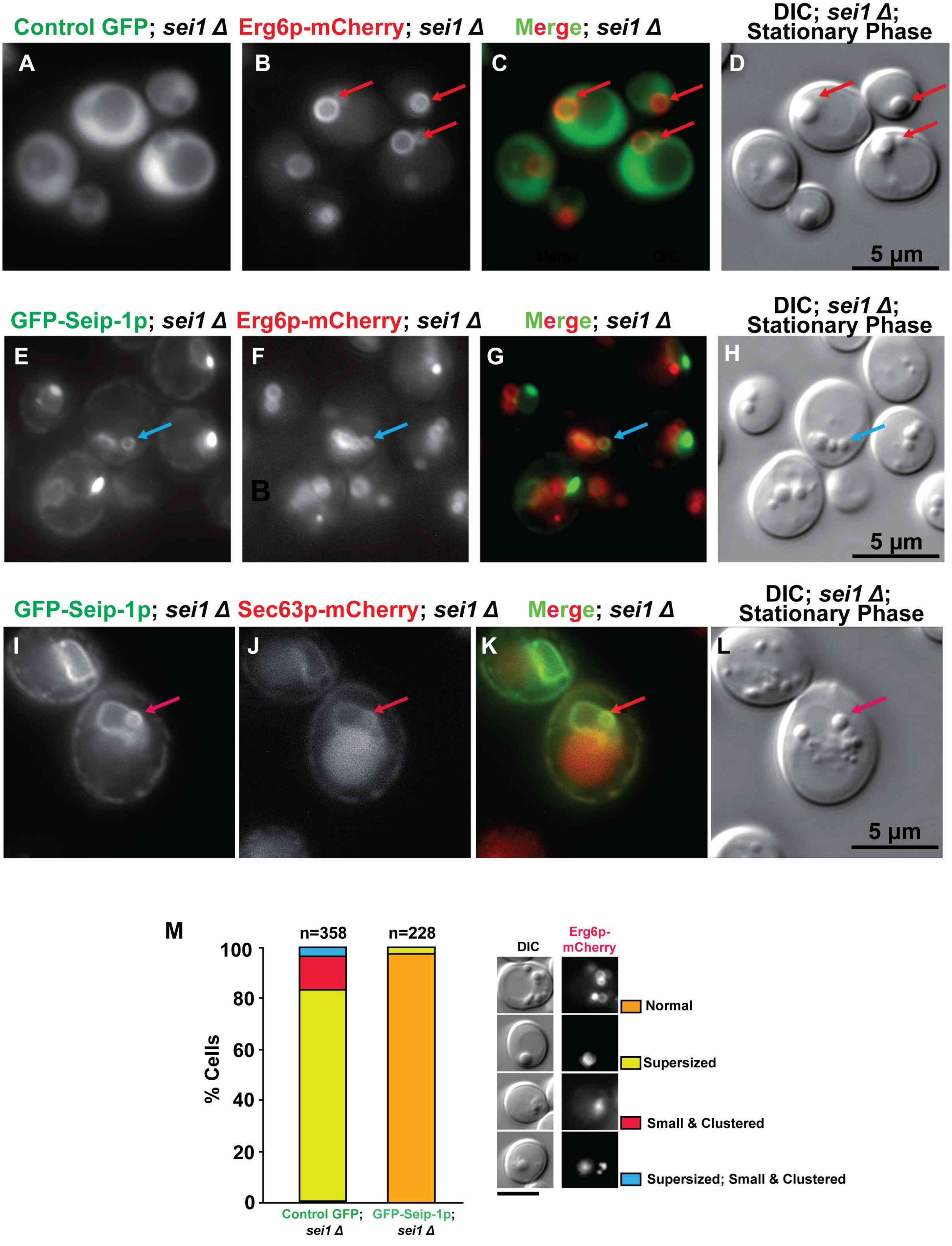

#### Supplemental Figure 2

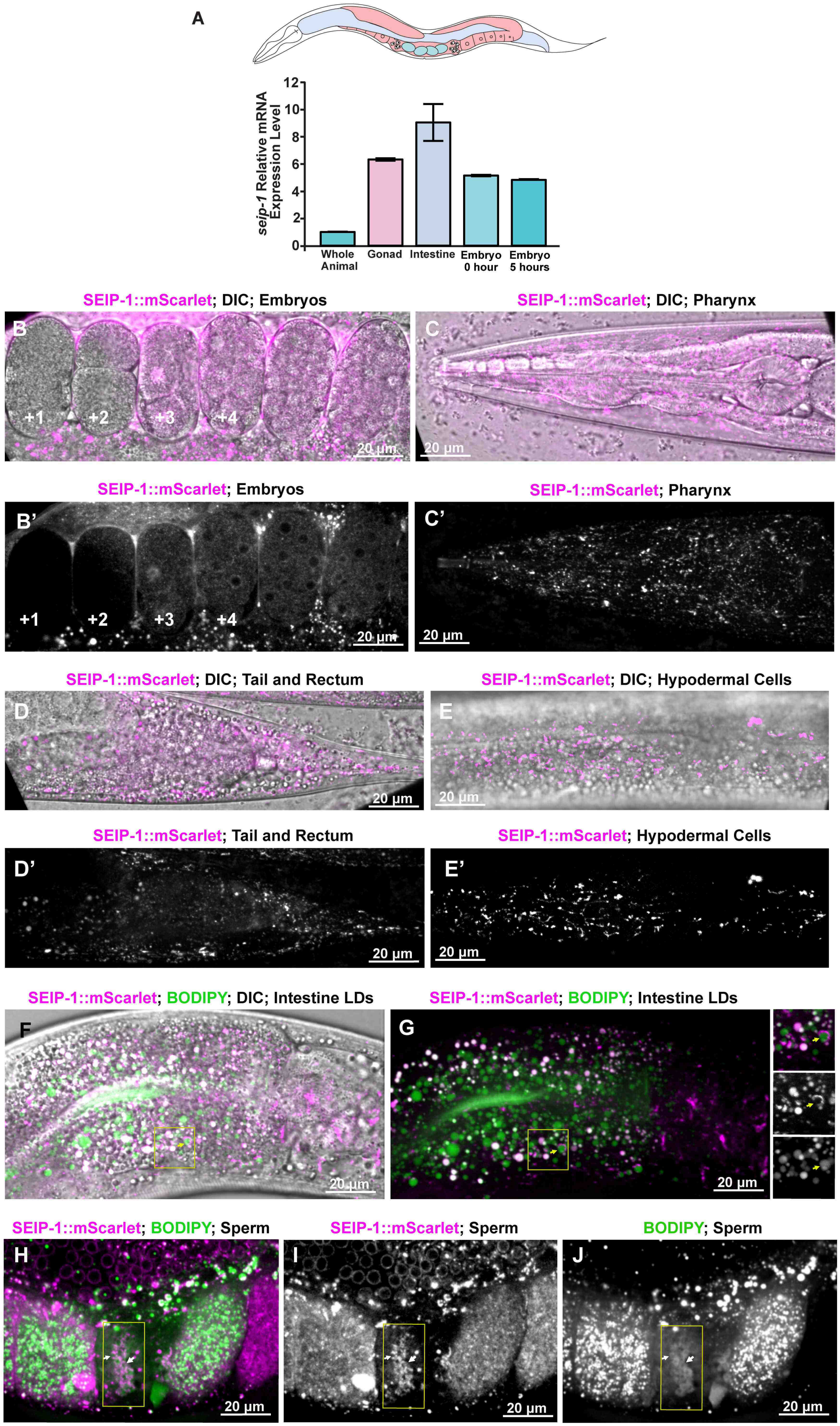

### Supplemental Figure 3

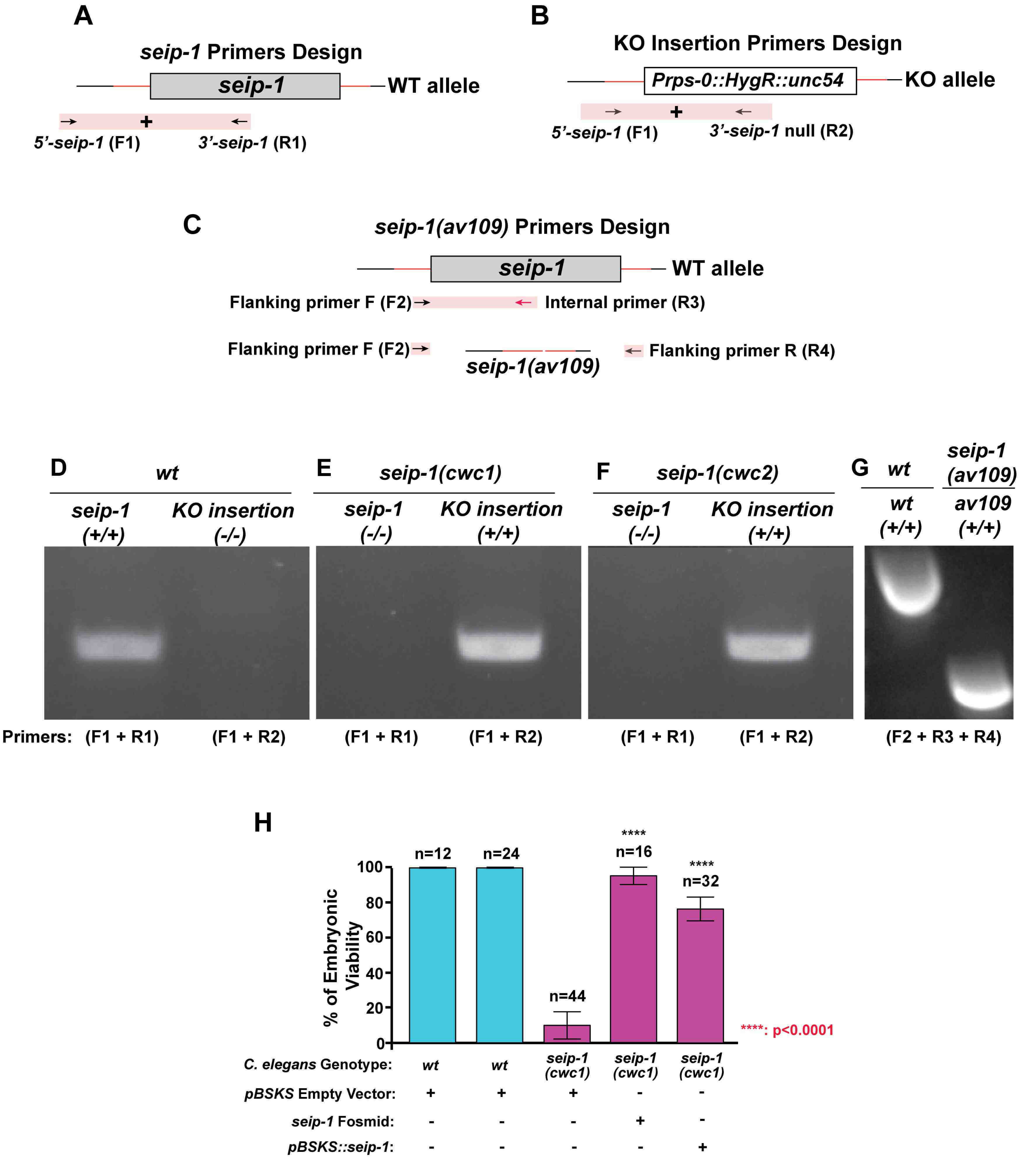

#### Supplemental Figure 4

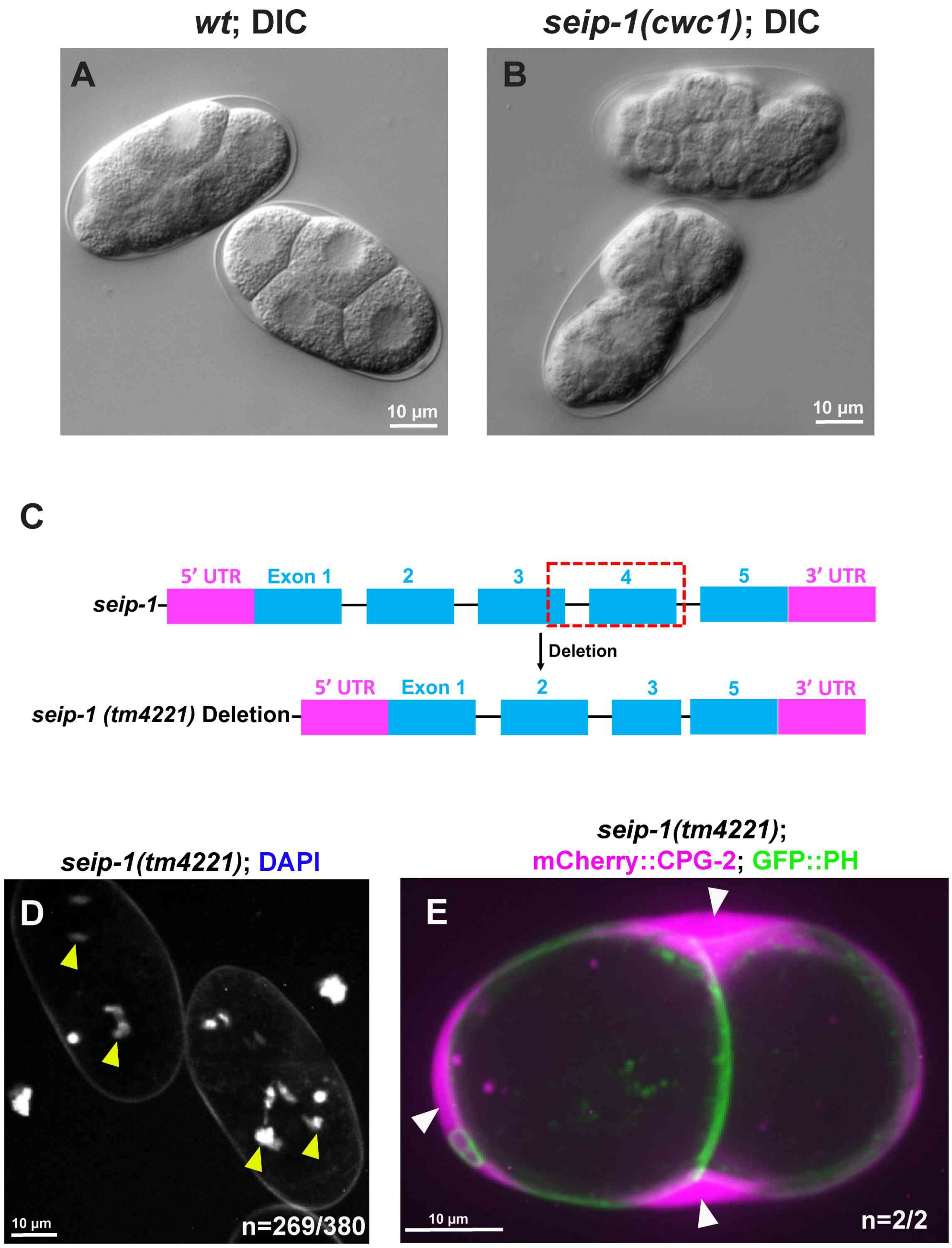

**Supplemental Figure 5**

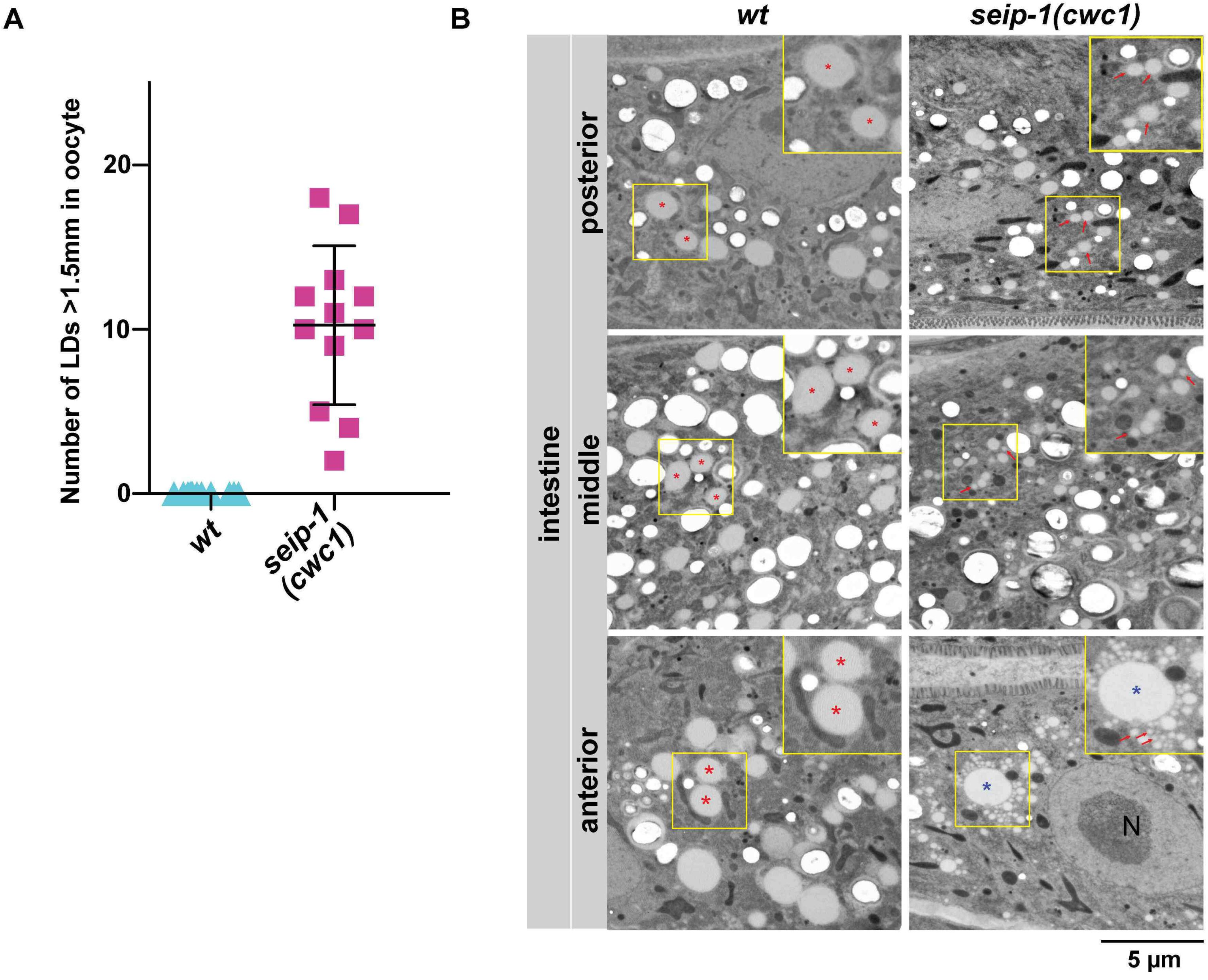

Supplemental Figure 6

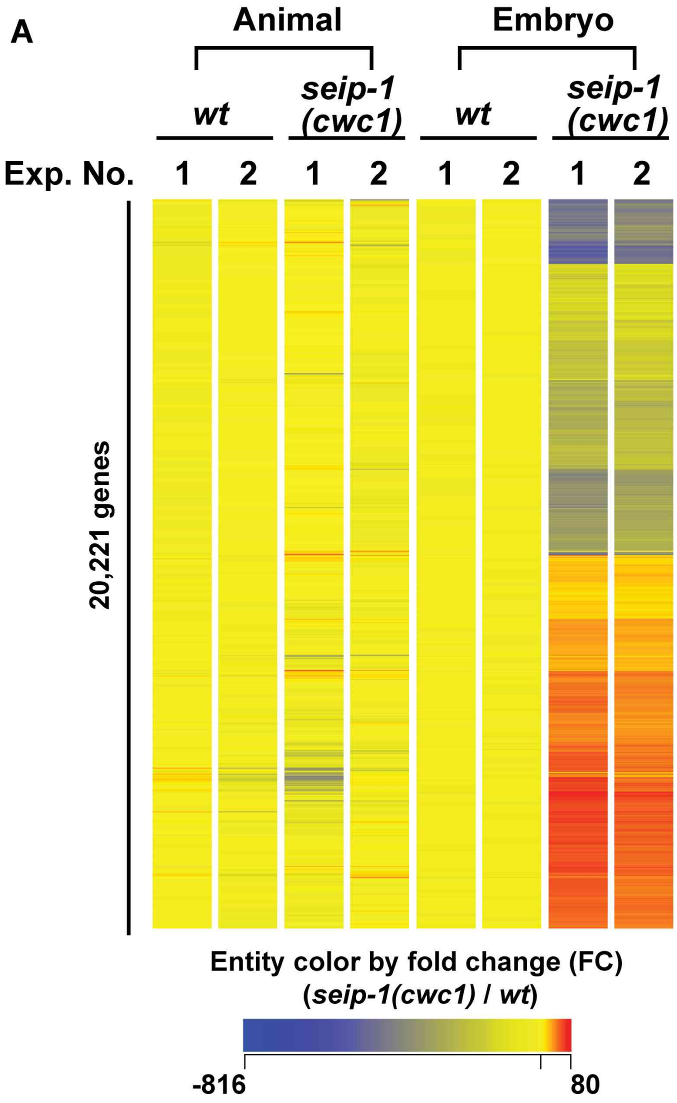


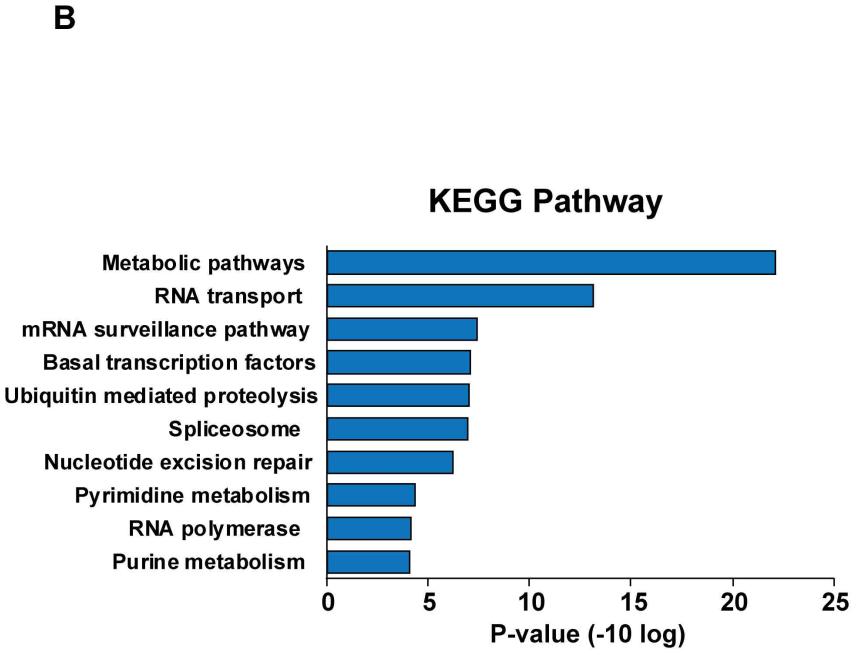

**Table 1 *C. elegans* and yeast strains list in the study.**

| No. Fig | Strain | Genotype |
|---|---|---|
| Fig.1 | AG444 | *seip-1(av169[seip-1::mScarlet]) V. CRISPR/Cas9 Edit* |
| | AG547 | *seip-1(av169[seip-1::mScarlet]) V; unc-119(ed3) III; ojIs23 [pie-1p::GFP::SP12 + unc-119(+)]* |
| | AG548 | *seip-1(av169[seip-1::mScarlet]) V; pwIs23 [vit-2::GFP]* |
| Fig.2 | N2 | Bristol (wild-type) |
| | AG363 | *seip-1(av109) V. CRISPR/Cas9 Edit. Deletion of coding region* |
| | CWC1 | *seip-1(cwc1) V. CRISPR/Cas9 Edit. seip-1 coding region replaced by HygR expression cassette.* |
| | CWC2 | *seip-1(cwc2) V. CRISPR/Cas9 Edit. seip-1 coding region replaced by HygR expression cassette.* |
| | OD344 | *unc-119(ed3) III; ItIs151 [pSO33; Pcpg-2::cpg-1SigSeq:: mCherry::cpg-2; unc-119(+)]; ltIs38 [pAA1; pie-1p::GFP::PH(PLC1δ1); unc-119(+)] III* |
| | AG549 | *seip-1(av109) V; unc-119(ed3) III; ItIs151 [pSO33; Pcpg-2::cpg-1SigSeq:: mCherry::cpg-2; unc-119(+)]; ltIs38 [pAA1; pie-1p::GFP::PH(PLC1δ1); unc-119(+)] III* |
| Fig.3 | N2 | Bristol (wild-type) |
| | CWC1 | *seip-1(cwc1) V. CRISPR/Cas9 Edit. seip-1 coding region replaced by HygR expression cassette.* |
| Fig.4 | N2 | Bristol (wild-type) |
| | AG363 | *seip-1(av109) V. CRISPR/Cas9 Edit. Deletion of coding region* |
| | CWC1 | *seip-1(cwc1) V. CRISPR/Cas9 Edit. seip-1 coding region replaced by HygR expression cassette.* |
| Fig. 5 | N2 | Bristol (wild-type) |
| | CWC1 | *seip-1(cwc1) V. CRISPR/Cas9 Edit. seip-1 coding region replaced by HygR expression cassette.* |
| Fig.6 | N2 | Bristol (wild-type) |
| | CWC1 | *seip-1(cwc1) V. CRISPR/Cas9 Edit. seip-1 coding region replaced by HygR expression cassette.* |
| Fig.7 | N2 | Bristol (wild-type) |
| | AG363 | *seip-1(av109) V. CRISPR/Cas9 Edit. Deletion of coding region* |
| | CWC1 | *seip-1(cwc1) V. CRISPR/Cas9 Edit. seip-1 coding region replaced by HygR expression cassette.* |
| | CWC2 | *seip-1(cwc2) V. CRISPR/Cas9 Edit. seip-1 coding region replaced by HygR expression cassette.* |
| | OD344 | *unc-119(ed3) III; ItIs151 [pSO33; Pcpg-2::cpg-1SigSeq:: mCherry::cpg-2; unc-119(+)]; ltIs38 [pAA1; pie-1p::GFP::PH(PLC1δ1); unc-119(+)] III* |
| | AG549 | *seip-1(av109) V; unc-119(ed3) III; ItIs151 [pSO33; Pcpg-2::cpg-1SigSeq:: mCherry::cpg-2; unc-119(+)]; ltIs38 [pAA1; pie-1p::GFP::PH(PLC1δ1); unc-119(+)] III* |
| Fig.8 | N2 | Bristol (wild-type) |
| | AG429 | *seip-1(av160[A185P]) V. CRISPR/Cas9 Edit.* |
| Fig.S1 | AG444 | *seip-1(av169[seip-1::mScarlet]) V. CRISPR/Cas9 Edit* |
| Fig.S2 | CWY3115 | *sei1Δ::HIS ERG6-mCherry::KAN* |

| | | |
|---|---|---|
| Fig.S3 | AG363 | *seip-1(av109) V. CRISPR/Cas9 Edit. Deletion of coding region* |
| | CWC1 | *seip-1(cwc1) V. CRISPR/Cas9 Edit. seip-1 coding region replaced by HygR expression cassette.* |
| | CWC2 | *seip-1(cwc2) V. CRISPR/Cas9 Edit. seip-1 coding region replaced by HygR expression cassette.* |
| Fig.S4 | N2 | Bristol (wild-type) |
| | CWC1 | *seip-1(cwc1) V. CRISPR/Cas9 Edit. seip-1 coding region replaced by HygR expression cassette.* |
| | FX14734 | *seip-1(tm4221)/hT1 V* |
| | OD344 | *unc-119(ed3) III; ItIs151 [pSO33; Pcpg-2::cpg-1SigSeq:: mCherry::cpg-2; unc-119(+)]; ltIs38 [pAA1; pie-1p::GFP::PH(PLC1δ1); unc-119(+)] III* |
| | AG549 | *seip-1(av109) V; unc-119(ed3) III; ItIs151 [pSO33; Pcpg-2::cpg-1SigSeq:: mCherry::cpg-2; unc-119(+)]; ltIs38 [pAA1; pie-1p::GFP::PH(PLC1δ1); unc-119(+)] III* |
| | AG550 | *seip-1(tm4221) V; unc-119(ed3) III; ItIs151 [pSO33; Pcpg-2::cpg-1SigSeq:: mCherry::cpg-2; unc-119(+)]; ltIs38 [pAA1; pie-1p::GFP::PH(PLC1δ1); unc-119(+)] III* |
| Fig.S5 | N2 | Bristol (wild-type) |
| | CWC1 | *seip-1(cwc1) V. CRISPR/Cas9 Edit. seip-1 coding region replaced by HygR expression cassette.* |
| Fig.S6 | N2 | Bristol (wild-type) |
| | CWC1 | *seip-1(cwc1) V. CRISPR/Cas9 Edit. seip-1 coding region replaced by HygR expression cassette.* |
| Fig.S7 | N2 | Bristol (wild-type) |
| | CWC1 | *seip-1(cwc1) V. CRISPR/Cas9 Edit. seip-1 coding region replaced by HygR expression cassette.* |
| Video.S1 | AG444 | *seip-1(av169[seip-1::mScarlet]) V. CRISPR/Cas9 Edit* |

#### Table 2 List of the sequence for the CRISPR design

| Strain | Genotype | Description | Sequence Name | Sequence 5'-3' | PAM |
|---|---|---|---|---|---|
| AG444 | *seip-1(av169[seip-1::mScarlet]) V* | Knock in mScarlet at C-terminus of *seip-1, mScarlet* was amplified from plasmid pMS050 | NEST1 crRNA | TTTTCTAAGATCCAAAAGTC | CGG |
| | | | Repair Template | AAAGAAGTGAAAAAAGAGATCAAAAAAGAAGAACC**A**GG**T**CT**C**TT**A**GA**C**CT**C**AG**G**AA**G**AG**G**AA**G**<br>GTCTCCAAGGGAGAGGCCGTCATCAAGGAGTTCATGCGTTTCAAGGTCCAAGCGCTCCGAGGGAC<br>GTCACTCCACCGGAGGAATGGACGAGCTCTACAAGTAGtgcttttcgattcaacatttaatactcaattt | |
| | | | Genotyping F1 | gaaaattccatccggcatcc | |
| | | | Genotyping R1 | ggcgctgacattaccacata | |
| | | | NEST2 crRNA | TTCAAGGTCCAAGCG**C**TCCG | |
| | | | Repair Template F1 | GCCGTCATCAAGGAGTTCATGCGTTTCAAGGTCCACATGGAGGGATCCATGAACG | |
| | | | Repair Template R1 | TAGAGCTCGTCCATTCCTCCGGTGGAGTGACGTCC**T**TC**T**GA**A**CGCTCGTATTGCTCGACGACGGTG | |
| AG363 | *seip-1(av109) V* | Deletion of coding region of *seip-1* | crRNA N-terminus | ccaaaagctacaatgccATG | AGG |
| | | | crRNA C-terminus | ACGAGCAGTCACGATTAACT | CGG |
| | | | Genotyping F1 | caaacttctctcgtggcacc | |
| | | | Genotyping R1 | ggcgctgacattaccacata | |
| | | | Genotyping Internal | gaaaattccatccggcatcc | |
| CWC1 and CWC2 | *seip-1(cwc1) V* and *seip-1(cwc2) V* | Knock out *seip-1* by replacing the *seip-1* coding sequence with *HygR* expression cassette. | crRNA N-terminus | ccaaaagctacaatgccATG | AGG |
| | | | crRNA C-terminus | aaagtagaaaatgaccagcg | AGG |
| | | | Repair Template | tggagcattttatttgacgcgcgatatctcttagcgaaaactacagtaagaacttttataacttaaataaaaaaatcgtttcaaatcggtaaaaattaccaatatcagtcag<br>aaaattaacttattttggttttctgattagttttctttcgatttttgaaaatgttatgaatttcaattgaaatttcatcgcccacgtttttaaataaatcgtttcataataaatcatgc<br>attcatcaatgcgagaccatggtctcgcattgatgaatgcatctacgctccaccgcactctccaaaatatttcatcgccttccggtgagtgtttgtgtgcacatctccgc<br>gtctctattcatctcataatctctgctctactaatttctctatttctcttctgcttttccatctcaatcatcgttctcttcgtcttcttcttcgtcttcttctcgtctttaatcactgttacc<br>gctcccatgacagttaattatattcaaacttctctcgtggcacccgcggctgcttcaaaaatgatgaaatattattttatattatcacatttattaacagactggaacggga<br>aaagttaccaaaagctacaatgcctggtcattttctactttgcctttataatttttataaacgatttttcgaatagattttcttaaactattgtgaatgtgctatattactggttgtt<br>cagccatgttttctatctgataatgtttatttttcaatttatttccccattattataaaatttattctttcagtttagtatgtggtaatgtcagcgccgccatgtcatcattgtctggc<br>cacgcgtccacaattcagccatcatgtctcggattctgcggtcgaaccgtcctcgtcggcaactattctgaagacgtggaggcgacgacgacggcggccggatcc<br>acgtcgttgtcaagatgtggaccatgttcatttggatatcgaaacaatgccatgtcaatatgcgaatcatgtgatactccactacaaccatacgattggatgtaccttttgt<br>ttatcgccttactcccattgcttctgcatatgcaatttatcagaatagccagaaagtagtaagtgattttcttcaataaaaacactgaataaattgtgtgactttaagtattag<br>gtattcaaaaaagacgtttctttagtattctatcgataaataaataaattaattcatttagaaaattgagccc | |
| | | | Genotyping F1 (*seip-1*) | acggcgttagtgctcgtgtcaaagg | |
| | | | Genotyping R1 (*seip-1*) | atgatgaactgacgacggaagaagg | |
| | | | Genotyping R2 (*HygR*) | agaagtactcgccgatagtggaaaccgacg | |
| AG429 | *seip-1(av160[A185P]) V* | Generate a point mutation A185P in *seip-1* | crRNA | ACGAGCAGTCACGATTAACT | CGG |
| | | | Repair Template | TATCAAGCCAAAACGAGCAGTCACGATTAACTCGG**GC**TCC**C**TCGATctggaaatagagaaataattagagttggac | |
| | | | Genotyping F1 | gaaaattccatccggcatcc | |
| | | | Genotyping R1 | ggcgctgacattaccacata | |

**Note:** Capital letters represent the ORF or exon sequence, small letters indicate the sequence from intron. Bolded letters indicate the optimized base needed for the CRISPR design.

#### Table 3 List of the sequence for the qPCR Primers

| Tested Gene | Sequence Name | Sequence 5'-3' | Tested Gene | Sequence Name | Sequence 5'-3' |
|---|---|---|---|---|---|
| *acox-1.1* | acox-1.1_F | CAAGTGGGCAAAGGAAAGTCC | *fasn-1* | fasn-1_F | TAAGCTGAAAAGTGTTCGCGGTA |
| | acox-1.1_R | ACTGACGGAAGAACATCTGTCTTGT | | fasn-1_R | CCAGCCCAGAGCCTATCCA |
| *acox-3* | acox-3_F | AAGATGGGTTTGCGATTTGG | *fat-1* | fat-1_F | AAGACCGCCGGAATCATG |
| | acox-3_R | GGATCCTTGTGATCTCTTTTGCA | | fat-1_R | CCTTTGCCTTCTCCTCGAGAGT |
| *act-1* | act-1_F | CTTCCCTCTCCACCTTCCAAC | *fat-2* | fat-2_F | CTTCACTACAACGTTACCCTCGACTA |
| | act-1_R | CGTCGTATTCTTGCTTGGAGATC | | fat-2_R | GACACCCTTTGCTTTATGAGTCAA |
| *cyp-31A2* | cyp-31A2_F | TCGTTCGCCCGGTCACT | *fat-3* | fat-3_F | CCACGTTGCAATCTGAATGC |
| | cyp-31A2_R | TGGGACGACGTCTGGTGAG | | fat-3_R | TTTGCACCATTCTTTCACATATTTC |
| *cyp-31A3* | cyp-31A3_F | GATTATCGTTCGCCCAGTCAC | *fat-4* | fat-4_F | GCACCATCTTTTCCCAACGA |
| | cyp-31A3_R | CGGCGTCTGGTAAGCTTCAT | | fat-4_R | TGGCATAACAGTGTTCAAGTTGTG |
| *daf-22* | daf-22_F | AATTGGTGGAGCTGGAGTAGTTG | *fat-5* | fat-5_F | GCAAGAAGTTCGGCTGTGAAA |
| | daf-22_R | GCTCCAGGGAATCCCAATCTA | | fat-5_R | TCCCAATTTGTGGAGCATTTT |
| *dgtr-1* | dgtr-1_F | CAATCTGTTCGAGGAGTACAAGCA | *fat-6* | fat-6_F | ATTATCGGCCGGCAGGTATC |
| | dgtr-1_R | TGAGTGTCGGGAGGAATGG | | fat-6_R | TTTTCCTCGTTGAATATCACATCC |
| *dhs-28* | dhs-28_F | TTCTTGAAAAGGCGAAGAAGTCA | *fat-7* | fat-7_F | GAGTTTATCAGCCGGCAGGTT |
| | dhs-28_R | ATTGAACGCTTCTGTCTGTTTACAA | | fat-7_R | TTTTCTTGATTCTTCACTTCCGTG |
| *elo-1* | elo-1_F | TCACCAATGCCAACTGTGATTT | *perm-1* | perm-1_F | TGGACCTTTTTCAACGCTACG |
| | elo-1_R | AAACTGCGAGCTTGAATACTGATG | | perm-1_R | TGGCTTGTATCCCAACATCAGA |
| *elo-2* | elo-2_F | CAAAAACGCTCACCAACCAA | *pod-2* | pod-2_F | TTGGAATCGGAGCCTACACG |
| | elo-2_R | CACAATGTTTATCTACTCCTGCTTGC | | pod-2_R | TGTGCTGAACGATTCGATGAG |
